# High radiosensitivity in the conifer Norway spruce (*Picea abies*) due to less comprehensive mobilisation of protection and repair responses compared to the radiotolerant *Arabidopsis thaliana*

**DOI:** 10.1101/2023.11.20.562501

**Authors:** Payel Bhattacharjee, Dajana Blagojevic, YeonKyeong Lee, Gareth B Gillard, Lars Grønvold, Torgeir Rhoden Hvidsten, Simen Rød Sandve, Ole Christian Lind, Brit Salbu, Dag Anders Brede, Jorunn E. Olsen

## Abstract

Risk assessment and protection of plant communities in contaminated ecosystems require in-depth understanding of differential sensitivity to chronic ionising radiation in plants. However, the contributing molecular factors to differential radiosensitivity among plant species are poorly understood. To shed light on this, we compared early events associated with protection, repair, and stress responses in gamma-irradiated (1-290 mGy h^−1^) seedlings of the radiosensitive conifer Norway spruce (*Picea abies*) and the radiotolerant *Arabidopsis thaliana*, by analysing growth, organelle and DNA damage, transcriptomes and the dynamics of antioxidant activities and expression of relevant genes. After 48 h of gamma radiation exposure, Norway spruce showed significantly reduced growth at 100-290 mGy h^−1^ and organelle damage, especially in mitochondria, at ≥ 1 mGy h^−1^ whereas *A. thaliana* showed normal vegetative growth at all dose rates, transiently delayed reproductive development at 290 mGy h^−1^ only, minor organelle damage only at ≥ 100 mGy h^−1^ and significantly less DNA damage than in Norway spruce at all dose rates. Comparative transcriptomics revealed that *A. thaliana* showed massive activation of genes related to DNA damage repair, antioxidants, and other stress responses at ≥ 1 mGy h^−1^ while Norway spruce mobilized transcription of such pathways only at ≥ 40 mGy h^−1^. The transcriptional activation of repair and protection responses at higher gamma dose-rates only and its absence in lower dose-rates, correlates with high radiosensitivity of Norway spruce, compared to the massive transcriptional activation from low dose-rates in the radiotolerant *A. thaliana*.

## 1. Introduction

Stress tolerance capacity in plants in general is highly variable and a defining factor for species niche and habitat preference. Ionising radiation is a stressor that can induce harmful effects to organisms due to damage to biological macromolecules such as DNA, proteins and lipids (Caplin and Wiley, 2018). Field experiments and surveys of radionuclide-contaminated plant communities and ecosystems have shown that species that are sensitive to chronic ionising radiation may succumb and be replaced by more tolerant species (Amiro and Sheppard, 1994; Caplin and Willey, 2018). Laboratory studies have confirmed that radiosensitivity of different plant species ranges over orders of magnitude (Caplin and Wiley, 2018; Blagojevic et al. 2019a and b; Duarte et al. 2023). Furthermore, previous studies have shown that conifers are among the most radiosensitive plant species whereas the herbaceous model plant *A. thaliana* is among the most radiotolerant ones (Sparrow and Miksche, 1961; Sparrow and Sparrow, 1965). Comparative studies employing gamma-irradiation of young seedlings under standardised conditions showed that the conifers Norway spruce and Scots pine (*Pinus sylvestris*) retained significantly more damage in terms of dose rate-dependent cellular damage including damage to the shoot apical meristem (SAM), DNA damage, inhibition of growth and even mortality at higher doses compared to the radiotolerant *A. thaliana* (Blagojevic et al. 2019a). However, the underlying mechanisms conferring tolerance or sensitivity to ionising radiation are not well understood. Such knowledge is needed to improve our ability to predict environmental consequences of nuclear accidents and release of radionuclides leading to exposure to high levels of ionising radiation.

The average natural background level of ionising radiation is estimated to 0.29 µGy h^−1^, while up to 15 µGy h^−1^ has been measured in areas with naturally elevated radionuclide levels (Caplin and Willey 2018). The nuclear power plant accident in Chernobyl in April 1986 as well as other accidents and experiments have shown that acute high doses (10-1000 Gy) can be lethal for plants and that radiosensitivity differs between species (UNSCEAR, 1996). In response to sub-lethal chronic radiation, adverse effects such as growth inhibition, reduced apical dominance (increased branching) and leaf chlorosis have been observed, particularly in conifers (Caplin and Wiley, 2018; Kashparova et al. 2020). A dose rate of 10 mGy d^−1^ (417 µGy h^−1^) has been suggested as a threshold value in the sense that lower dose rates have “no detrimental effects for populations of terrestrial plants in the field” (International Atomic Energy Agency [IAEA], 1992; Caplin and Willey 2018; UNSCEAR). However, this does not appear to hold true for boreal conifer forests. In a long-term field irradiation experiment in

Canada persistent effects on forests including conifer trees were observed at chronic dose rates ≥ 100 µGy h^−1^ (Amiro and Sheppard 1994). In addition to the differential radiosensitivity between species (Blagojevic et al. 2019a; Vanhoudt et al. 2014; Choi et al. 2019), effects of ionising radiation depend on dose rate, duration (acute or chronic), total dose, life stages and the co-occurrence of other contaminants and environmental conditions (Garnier-Laplace et al. 2013).

The sensitivity of plant species to acute irradiation has been suggested to be attributed to the efficiency of DNA repair systems and genome organisation and size (De Micco et al. 2011). *A. thaliana* has a small genome of 135 Mb, whereas conifers like Norway spruce and Scots pine have giant genomes of 20 and 23 Gb (Nystedt et al. 2013; Zimin et al. 2017). On the other hand, the sensitivity to chronic irradiation appears to be regulated by delicate mechanisms on different levels of biological organisation (Caplin and Willey 2018; Kim et al. 2019). However, factors that define sensitivity to chronic exposure are poorly understood and apparently have functional rather than structural origin, depending on the molecular architecture of stress-responsive pathways of different plant species.

Ionising radiation can cause damage both directly and indirectly via the formation of free radicals and subsequent reactive oxygen species (ROS) (Caplin and Willey 2018). As one of the key forces of adaptive responses, non-lethal levels of ROS can modulate cellular responses related to redox homeostasis and initiate retrograde signalling between organelles, such as mitochondria, chloroplasts and the endoplasmic reticulum, and the cell nucleus, thereby affecting gene expression (Volkova et al. 2022). Plants possess antioxidant systems involving enzymes such as catalases (CAT), superoxide dismutase (SOD) and peroxidases (PER) as well as metabolites such as ascorbate, phenolic compounds and glutathione (GSH) that can neutralise ROS and thereby prevent oxidative damage (Dumanović et al. 2020). Studies of specific plant species including *A. thaliana*, have revealed induction of antioxidant-related genes, including *SOD* and *ASCORBATE PEROXIDASE* (*APX*) following ionising radiation exposure (van de Walle et al. 2016; Kim et al. 2011) but there is limited information about antioxidant homeostasis in conifers.

DNA strand breaks induced by ROS or directly by ionising radiation trigger the activation of DNA damage repair (DDR) pathways (Kim et al. 2019). A range of homologous-recombination (HR) repair-mediated DDR genes known to be activated by the plant-specific transcription factor *SUPRESSOR OF GAMMA RESPONSE 1 (SOG1)* were significantly upregulated by gamma irradiation at 100 Gy in *A. thaliana* (Bourbousse et al. 2018). The initiation of the DDR is closely interlinked with cell cycle stalling through the activation of cell cycle checkpoints by regulating cyclin (CYC)-cyclin-dependent kinase (CDK)-complexes (Qi and Zhang 2019), further leading to DNA damage-induced endoreduplication (Nisa et al. 2019). In addition, transcriptional recruitment of DDR proteins in response to gamma radiation requires chromatin remodelers involved in DNA methylation and histone 2A (H2A) modification (Choi et al. 2019; Kim 2019; Kim et al. 2021). However, there is a lack of knowledge related to molecular events associated with DDR and other relevant pathways in conifers exposed to ionising radiation.

Knowledge of the factors determining the radiosensitivity of plant species is crucial for risk assessment and the protection of ecosystems from ionising radiation hazard. Although transcriptional regulation in response to very high doses of ionising radiation has been investigated, albeit to a limited extent, in *A. thaliana* and a few other species (Choi et al. 2021; Vanhoudt et al. 2014), the mechanisms leading to the differential radiosensitivity among plant species are poorly understood. Hence, this study aimed to provide an overview of mechanisms involved in plant radiosensitivity by employing gamma irradiation under standardised conditions and concomitant analyses of growth, organelle- and DNA damage as well as transcriptomics and dynamics of antioxidant activities and expression of repair- and protection-related genes. In this respect, early physiological, cellular, and molecular effects were compared in young seedlings of the radiosensitive conifer Norway spruce and the radiotolerant *A. thaliana* in order to provide novel insights into mechanisms underlying their differential radiosensitivity.

## 2. Materials and methods

### 2.1. Plant materials and pre-growing conditions

Seeds of the Landsberg erecta (L*er*) accession of *A. thaliana* (L) Heyhn, and Norway spruce (*Picea abies* L. H. Karst) from the provenance CØ1 from Halden, Norway (59°N latitude, Skogfrøverket, Hamar, Norway) were surface-sterilised and sowed according to the protocol in (Blagojevic et al. 2019a). The seeds were germinated for 6 days at 20°C under a 16 h photoperiod with a photosynthetic photon flux density (PPFD) of 30 μmol m^−2^ s^−1^ at 400-700 nm (TL-D 58W/840 lamps, Philips, Eindhoven, The Netherlands), as measured at the top of the Petri dishes with a Li-Cor Quantum/Radiometer/Photometer (LI-250, LI-COR, USA).

### 2.2. Gamma irradiation of seedlings and growing conditions during the exposure

Six days old seedlings of *A. thaliana* and Norway spruce were exposed for 48 h using the FIGARO gamma irradiation facility (^60^Co; 1173.2 and 1332.5 keV γ-rays) at the Norwegian University of Life Sciences (Lind et al. 2019)). At day 6-8 of germination, the two species are at as similar a developmental stage as possible, prior to epicotyl expansion in either species under the environmental conditions used. Petri dishes (3.5 cm diameter) with seedlings were placed at different distances from the gamma source to obtain the average dose rates to water 1, 10, 20, 40, 100 and 290 mGy h^−1^ (Table 1). These 48-h irradiation experiments were repeated three times (Table S1. For one of these experiments, the highest dose rate was 250 mGy h^−1^ as 290 mGy h^−1^ could not be obtained due to the decay of the ^60^Co source. In additional experiments aiming at studies of dynamic effects early during gamma irradiation on gene expression and antioxidant contents, seedlings of both species were exposed to 0, 1, 10, 20, 40 and 100 mGy h^−1^ for 12, 24 or 48 h, with 2 repeated experiments for each duration (Table S1). The dose rates to water and dose rate intervals (in front and back of the petri dishes) were calculated from the air kerma rates according to (Hansen et al. 2019) as previously described (Blagojevic et al. 2019b). The total doses were calculated by multiplying the absorbed dose rates to water and exposure duration. During the exposure the room temperature was 20°C and the photoperiod 12 h with an irradiance of 55 μmol m^−2^ s^−1^ from high pressure metal halide lamps (HPI-T Plus 250W lamps, Philips, The Netherlands). The red:far red (R:FR) ratio was measured to 3.5 using a 660/730 nm Skye sensor (Skye Instruments, UK).

**Table 1.** The gamma radiation dose rates and total doses applied in experiments with 12, 24 and 48 h exposure of seedlings of *Arabidopsis thaliana* and Norway spruce from day 7 after sowing, using a ^60^Co source.

| Dose rate<br>(mGy h<sup>-1</sup>) | Dose rate<br>interval<br>(mGy h<sup>-1</sup>) |  | Total<br>dose (Gy)<br>after 48 h<br>exposure | Total dose<br>interval (Gy)<br>after 48 h<br>exposure |  | Total dose<br>(Gy) after<br>24 h<br>exposure | Total dose<br>interval (Gy) after<br>24 h exposure |  | Total dose<br>(Gy) after 12<br>h exposure | Total dose<br>interval (Gy)<br>after 12 h<br>exposure |  |
| --- | --- | --- | --- | --- | --- | --- | --- | --- | --- | --- | --- |
|  | Min | Max |  | Min | Max |  | Min | Max |  | Min | Max |
| 290 | 255 | 325 | 13.9 | 12.24 | 15.60 | 7.0 | 6.12 | 7.80 | 3.5 | 3.06 | 3.90 |
| 100 | 88 | 112 | 4.8 | 4.22 | 5.38 | 2.4 | 2.11 | 2.69 | 1.2 | 1.06 | 1.34 |
| 40 | 35 | 45 | 1.9 | 1.68 | 2.16 | 1.0 | 0.84 | 1.08 | 0.48 | 0.42 | 0.54 |
| 10 | 8,8 | 11,2 | 0.48 | 0.42 | 0.54 | 0.24 | 0.21 | 0.27 | 0.12 | 0.11 | 0.13 |
| 1 | 0,88 | 1,12 | 0.048 | 0.042 | 0.054 | 0.024 | 0.021 | 0.027 | 0.012 | 0.011 | 0.013 |
| 0 | - | - | 0 | - | - | 0 | - | - | 0 | - | - |

### 2.3. Post-irradiation growing conditions

The post-irradiation effects of the 48 h of gamma exposure were recorded after transfer of seedlings to pots filled with S-soil (Hasselfors Garden AS, Örebro, Sweden), one plant per pot. All plants were kept in growth chambers (manufactured by Norwegian University of Life Sciences) with temperature, relative air humidity, R:FR ratio, and photoperiod maintained as previously described (Blagojevic et al. 2019a).

### 2.4. Growth parameter recordings

At the end of the gamma irradiation, seedlings from both species were placed between two transparent plastic sheets with a millimetre paper on top and scanned. The hypocotyl and radicle lengths of Norway spruce and the lengths of the hypocotyl and cotyledons in *A. thaliana* were measured using Image J (imagej.nih.gov) with 28-35 and 29-40 seedlings per gamma dose rate for the respective species.

Post-irradiation, the number of rosette leaves as well as the number of plants with floral buds, inflorescence and open flower buds were recorded in 23-24 *A. thaliana* plants per gamma dose rate in time courses up to 75 days post-irradiation (dpi), and percentages of plants with floral buds, inflorescence and open flowers were calculated. Leaf width and leaf length were measured at 34 and 41 and height of the inflorescence at 64 dpi. In Norway spruce, shoot elongation, the number of needles and shoot diameter were recorded in a time course of 59 dpi in 11-22 plants per dose rate. The shoot diameter was calculated by averaging two perpendicular measurements from needle tip to needle tip across the plant at the shoot apex. Also, plant height was measured from the pot edge to the shoot apical meristem and the cumulative shoot elongation was calculated.

### 2.5. Histological and cytological studies of shoot tips

Histological studies of shoot apical meristems (SAMs) at the end of the gamma irradiation were performed according to (Lee et al. 2017). Shoot tips of Norway spruce and *A. thaliana* were then fixed in 4 % paraformaldehyde and 0.025 % glutaraldehyde in phosphate buffered saline (PBS, pH 7.0), vacuum infiltrated at room temperature and stored at 4°C. The fixed tissues were washed in PBS solution and dehydrated in a graded ethanol solution. Approximately 3 mm of the shoot tips of each of 5 plants per species per treatment were infiltrated and embedded in LR White (London Resin Company, UK). After polymerisation, 1 µm sections were made using an Ultracut Leica EM UC6 microtome (Leica, Germany), stained with toluidine blue O and inspected using a light microscope with bright field optics (DM6B, Leica).

For cytological studies, LR white-embedded samples from 5 plants per species per treatment were sectioned into 70 nm ultrathin sections, (Leica EMUC6), stained with uranyl acetate and lead citrate before examining by transmission electron microscopy (TEM) (FEI Morgagni 268, United States).

### 2.6. Comet assay for analysis of DNA damage

Single and double-strand DNA breaks at the end of the gamma irradiation and 77 dpi were quantified using the comet assay (Gichner 2003) with some modifications (Blagojevic et al. 2019a). Briefly, this included isolation of cell nuclei and electrophoresis under alkaline pH, resulting in a comet-like appearance of damaged DNA due to movement out of the nuclei. This was followed by quantification of the DNA in the comet tail using fluorescence microscopy. Three biological replicates per dose rate, each consisting of 3-4 mm of shoot tips from 3 plants of Norway spruce or 5 plants of *A. thaliana*, were used to assess DNA damage. For each sample, about 180-200 cell nuclei were scored, with 60-70 nuclei scored in each of 3 technical replicates (gels) and the median value was calculated.

### 2.7. RNA isolation for RNA sequencing

At the termination of the 48-h gamma irradiation, four biological replicates per gamma dose rate (0, 1, 10, 40, 100 (both species) and 290 mGy h^−1^ (*A. thaliana* only due to lack of growth inhibition, organelle and DNA damage at lower dose rates)) were harvested in liquid nitrogen for subsequent RNA isolation. Due to the differences in seedling size between the two species, different numbers of shoots were used to obtain approximately 50 mg tissue per sample. It was also not practically feasible to use the same number of shoots per sample for the relatively large Norway spruce seedlings as for *A. thaliana,* due to space limitations in the gamma radiation sector at the highest dose rates. For *A. thaliana*, total RNA was extracted from 20-30 shoots per sample using the RNeasy Plant Mini Kit (QIAGEN Inc., Valencia, CA) following the manufacturer’s protocol. For Norway spruce total RNA was extracted from 4 shoots per sample using the Masterpure Complete DNA and RNA Purification Kit (Epicenter, Madison, USA) following the manufacturer’s protocol, except that 3 µl beta-mercaptoethanol per sample replaced 1,4-dithiothreitol (DTT) and 0.5% polyvinylpyrrolidone (PVP, mw 360 000, Sigma Aldrich, Germany) was added to the extraction buffer. The purified RNA was quantified using a NanoDrop ND-1000 Spectrophotometer (Thermo Scientific, USA) and the quality assessed with an Agilent 2100 Bioanalyzer with an RNA 144000 Nano Kit (Agilent technologies, USA).

For each species, the results for each gamma radiation dose rate were normalised to its own control group in the RNA sequencing and qPCR analyses described in section 2.8-2.10. The analyses thus compare the relative changes within each species rather than absolute values between species. Thus, the different number of shoots per sample for the two species is not likely to affect the results. Also, the use of fewer seedlings in the Norway spruce samples do not necessarily imply lower genetic variation compared to the *A. thaliana* samples. The inbred *A. thaliana* L*er* accession (a laboratory strain; Aloso-Blanco and Koorneef 2000) likely exhibits less genetic variation than the Norway spruce provenance, whose seeds were collected from natural populations.

### 2.8. RNA sequencing

The transcriptomes of the shoots were sequenced with two repeated lanes per sample, on an Illumina HiSeq4000 platform at the Norwegian Sequencing Centre (Oslo, Norway) using the Strand specific 20xTruseq RNA library preparation (paired end, read length 150 bp) as recommended by the manufacturer (Illumina, San Diego, CA, USA). The raw sequence data were submitted to the ArrayExpress under the accession number E-MTAB-8081 for Norway spruce and E-MTAB-11120 for *A. thaliana*.

### 2.9. Differential gene expression analyses and functional term enrichment

The transcript abundances were estimated using the Salmon program (version 0.10.2) (Patro et al. 2017), comparing with the genomes of either Norway spruce (*Picea abies* 1.0 assembly, plantgenie.org) (Nystedt et al. 2013) or *A. thaliana* (TAIR10, arabidopsis.org) (Berardini et al. 2015). Gene expression was compared between the samples at different gamma radiation dose rates and the unexposed control samples using the DESeq2 (version 1.34.0) (Love et al. 2014). Genes were classified as differentially expressed genes (DEGs) if the False Discovery Rate (Benjamini and Hochberg, correction) adjusted p-values were < 0.05.

Gene Ontology (GO) annotations of Norway spruce genes, as well as spruce to *A. thaliana* orthologs by best sequence matches, were downloaded from plantgenie.org. The *A. thaliana* orthologs were used to associate the Norway spruce genes with KEGG pathways. Thus, assumptions of gene orthologs in the two species are based on sequence similarity since validation of gene function is, with few exceptions, lacking in Norway spruce. Enrichment of GO or KEGG terms in sets of DEGs was found using topGO (version 2.34.0) or kegga from the limma package (version 3.38.2), respectively.

### 2.10. RT-qPCR analysis of dynamics in gene expression under ionising radiation

Based on the significant regulation in one or both species in response to 48 h gamma irradiation, 11 DNA repair- and cell cycle control genes as well as 4 antioxidant- and 2 retrograde signalling-related genes were selected for RT-qPCR analysis of their dynamic regulation after 12, 24 and 48 h of gamma irradiation at 0, 1, 10, 20, 40 and 100 mGy h^−1^ (Table S2). RNA was isolated as described above from four biological replicates per dose rate, each consisting of shoots from eight and 20-30 seedlings for Norway spruce and *A. thaliana*, respectively. cDNA was synthesised using SuperScript™ IV VILO™ cDNA Synthesis Kit (Invitrogen, Carlsbad, CA, US) following the manufacturer’s protocol, and no enzyme controls (-RT) were made for each sample.

Gene sequences were retrieved from NCBI (ncbi.nlm.nih.gov/gene) and TAIR10 (arabidopsis.org) for *A. thaliana* and plantgenie.org for Norway spruce and exon-exon-spanning primers were designed to avoid non-specific amplification from any genomic DNA contamination in the RNA samples, using the online tools Oligoanalyzer (eu.idtdna.com/calc/analyzer) and Primer Blast (ncbi.nlm.nih.gov/tools/primer-blast) with default parameters. Amplifications were performed in a 7500 Fast Real-time PCR system (Applied Biosystems, Foster city, USA) using Platinum Quantitative PCR Supermix-UDG and SYBRGreen (Thermo Fisher, Carlsbad, CA, US), using standard protocol.

To compare the relative transcript levels of the selected genes in the samples exposed to different gamma radiation dose rates at different time points with the unexposed controls (0 mGy h^−1^), the comparative ΔΔCT method was used (Livak and Schmittgen 2001). The expression of the target genes was normalised by the mean expression of the three reference genes, *ACTIN* (*ACT*), *ELONGATION FACTOR 1*α (*EF-1*α) for both species and *UBIQUITIN* and α*-Tubulin* for *A. thaliana* and Norway spruce respectively.

### 2.11. Antioxidant assays

Shoot samples for analyses of PER and CAT concentration, SOD activity and flavonoid content in seedling shoots of both plant species were flash-frozen in liquid nitrogen after 12, 24 and 48 h of gamma irradiation at 0, 1, 10, 20, 40 and 100 mGy h^−1^. Samples of 50 mg -100 mg were homogenized maintaining freezing conditions with stainless steel beads in a tissue homogenizer (Retsch, Haan, Düsseldorf, Germany). The samples were then mixed with appropriate buffers according to the protocol for each of the different antioxidant kits used (OxiSelect, Cell Biolabs. Inc., San Diego, CA, USA). The mixtures were vortexed and subsequently centrifuged to precipitate cellular debris, and the clear lysates were collected for the assays. The assays were performed in room temperature following the instructions in the manufacturers’ protocol for each antioxidant. Colorimetric analyses were done in a plate reader (Microplate reader, Biochrom (ASYS UVB340)). For each antioxidant, three (peroxidase, catalase, SOD) or four (flavonoid) biological replicates, each consisting of materials from 20-30 shoots for *A. thaliana* and 4 shoots for Norway spruce, were analysed per dose rate and gamma irradiation duration with 2-3 technical replicates of each.

### 2.12. Statistical analysis for growth recordings, DNA damage, qPCR and antioxidant assay

DNA damage, hypocotyl, cotyledon and shoot length, as well as post-irradiation growth parameters at the final sampling time (34, 41 or 64 dpi for *A. thaliana* and 59 dpi for Norway spruce) were tested for normality and homoscedasticity by the Ryan-Joiner and Levene tests, respectively. One-way ANOVA tests (glm model in Minitab 21.1, Minitab Inc, PA, USA) were then used to test for differences in these response variables between plants exposed to 48 h of gamma radiation versus unexposed control plants. Due to the lack of normality and homoscedasticity of the non-transformed RT-qPCR gene expression and antioxidant analyses data, log-transformed values were used in two-way ANOVA for individual species (dose rate and gamma-irradiation duration as factors) and three-way ANOVA for comparison of species (plant species, dose rate and gamma-irradiation duration as factors). Tukey’s post hoc test was used to test for differences between means (p ≤ 0.05).

## 3. Results

### 3.1. Gamma irradiation for 48 h affects growth more negatively in Norway spruce but not A. thaliana

In *A. thaliana*, no significant effect of the 48-h of gamma irradiation on hypocotyl and cotyledon lengths was identified (Fig. 1A, B). Also, the number of rosette leaves developing post-irradiation was not significantly affected (Fig. 1C). Only a slight, significant reduction in the width and length of rosette leaves was observed at the highest dose rate compared to the unexposed control at 34 and 41 dpi, but apart from this, there was no clear dose response-relationship (Fig. S1A, B). Although post-irradiation flowering was slightly delayed at 250-290 mGy h^−1^, eventually all plants flowered (63 dpi) and showed very similar overall morphology, flowering, and silique development (77 dpi) (Fig. 1C, S1D, E, F, G). In Norway spruce, the hypocotyl length was significantly reduced (27%) after 48 h of gamma irradiation at 290 mGy h^−1^, but the radicle length was not affected (Fig. 1D, E). At 59 dpi, the shoot diameter and number of needles were significantly reduced by 5% and 12%, at 100 mGy h^−1^ and 9% and 35% at 290 mGy h^−1^ whereas the cumulative growth did not differ significantly between the gamma dose rates but (Fig. 1F, G, H, Fig. S2). However, the growth inhibition effects were no longer as evident at 77 dpi (Fig. 1H).

**Fig. 1.**
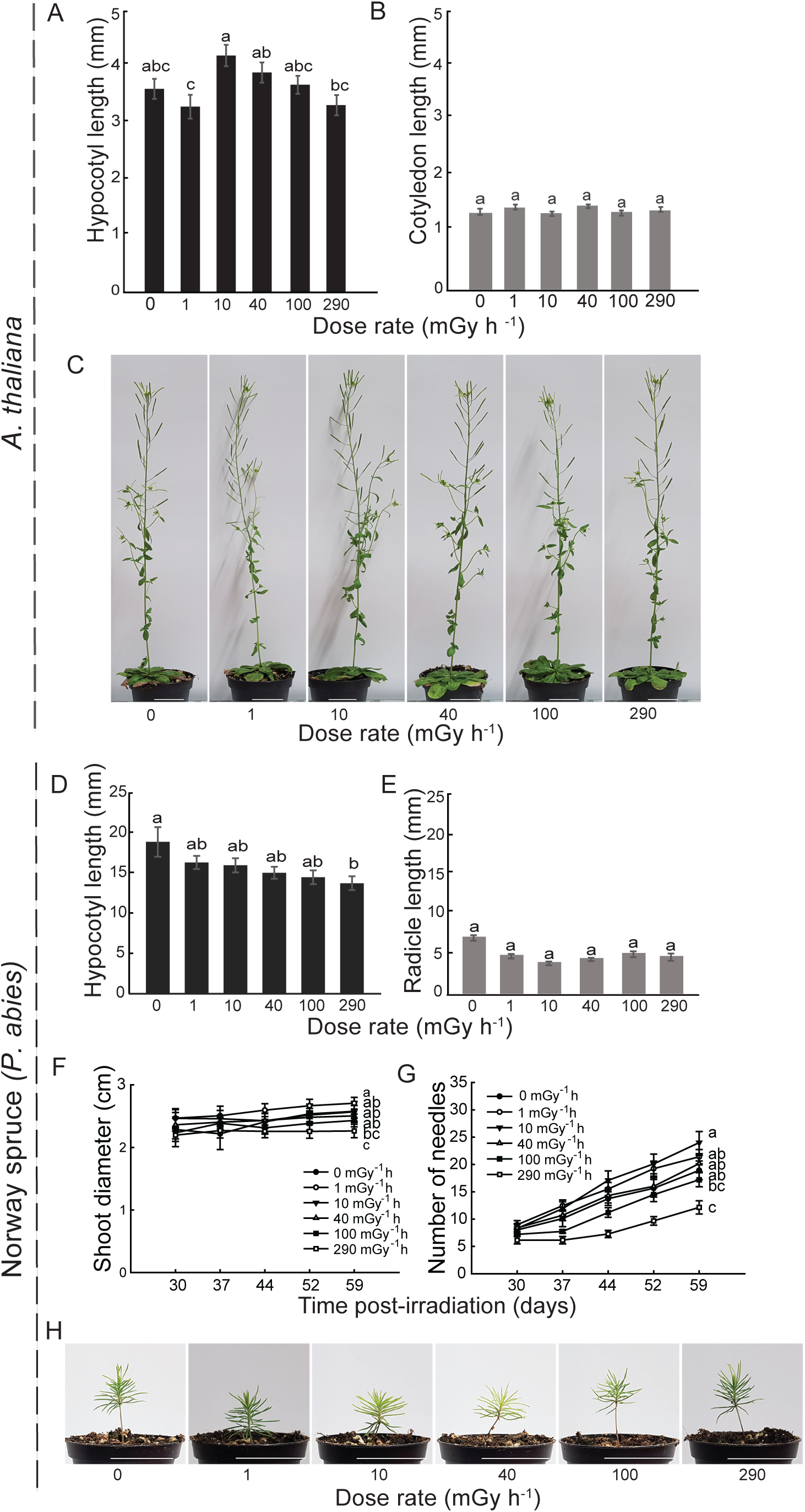
Effect of 48 h of gamma irradiation from day 7 after sowing on growth in *A. thaliana* and Norway spruce (*P. abies*). (a) Hypocotyl and (b) cotyledon length in *A. thaliana* (n = 29-40) after the 48-h irradiation and (c) morphology 77 days post-irradiation (dpi). (d) Hypocotyl and (e) radicle length of Norway spruce (n = 28-35) after the 48-h irradiation and (f) shoot diameter and (g) number of needles in time courses (n = 11-22) and (h) morphology 77 dpi. Scale bars: 3 cm. Means ± standard errors are shown. Different letters within each figure denotes significant difference (p ≤ 0.05) based on analysis of variance followed by Tukey’s test.

### 3.2. More severe gamma radiation-induced organelle damage in Norway spruce than A. thaliana

Normal dome-shaped shoot apical meristems with compact cells were observed in both species after 48 h of gamma irradiation (Fig. 2). In *A. thaliana*, slightly damaged mitochondria were observed at 100-290 mGy h^−1^, but the chloroplasts and cell nuclei appeared normal at all dose rates compared to the unexposed control except that a few cells showed less dense nuclei at ≥ 1 mGy h^−1^ (Fig. 3A, B, C). In comparison, in Norway spruce, disrupted nuclear and mitochondrial membranes were observed at ≥ 1 mGy h^−1^, as well as swollen chloroplasts with large starch bodies and disrupted thylakoid membranes at ≥ 40 mGy h^−1^ (Fig. 3D, E, F).

**Fig. 2.**
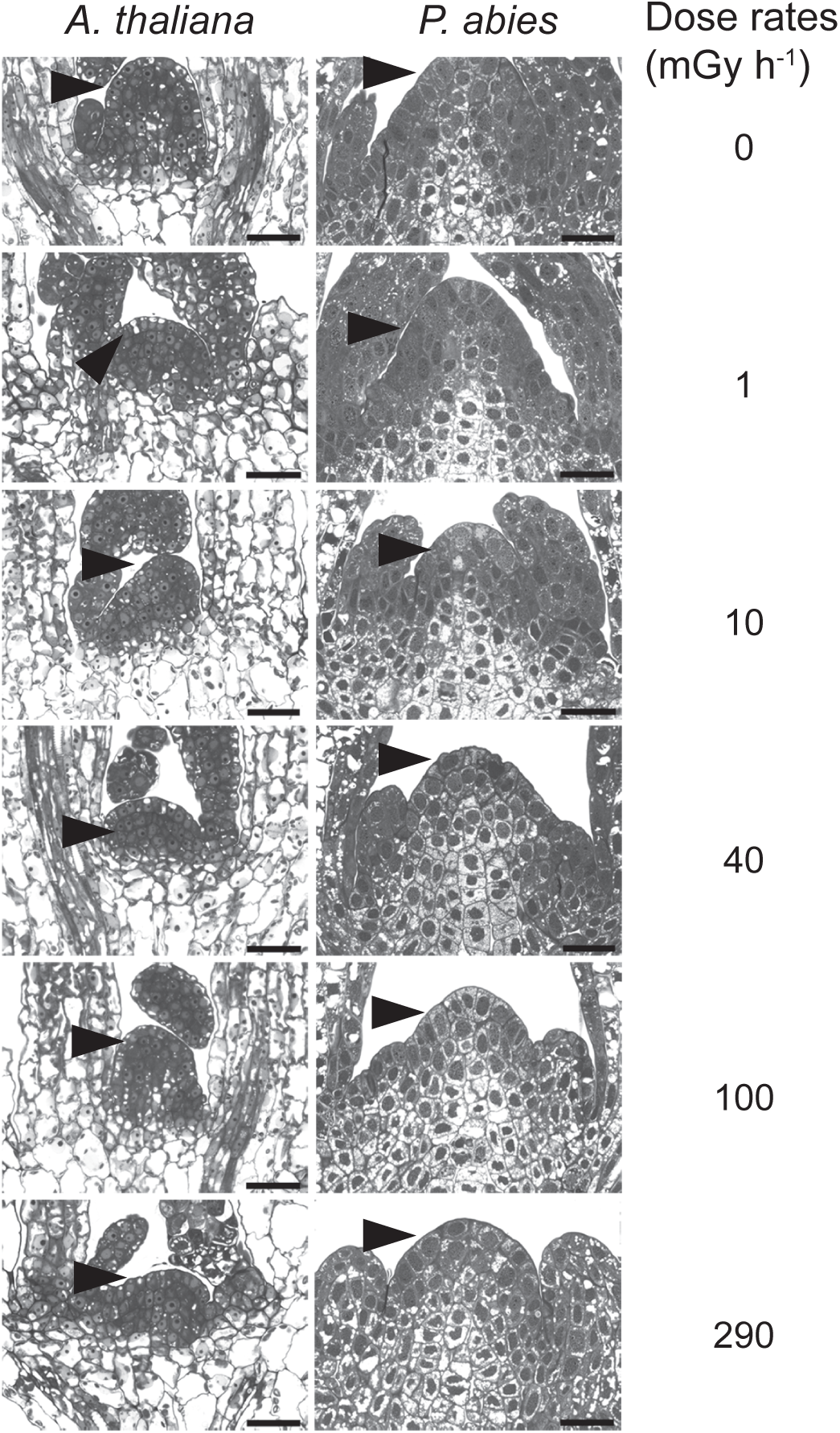
Light micrographs of shoot tips with the shoot apical meristem of *A. thaliana* and Norway spruce (*P. abies*) after 48 h of gamma irradiation from day 7 after sowing. The black arrowhead in each subfigure points to the apical dome of the SAM. Five plants were investigated per treatment for each species. Scale bars: 25 µm.

**Fig. 3.**
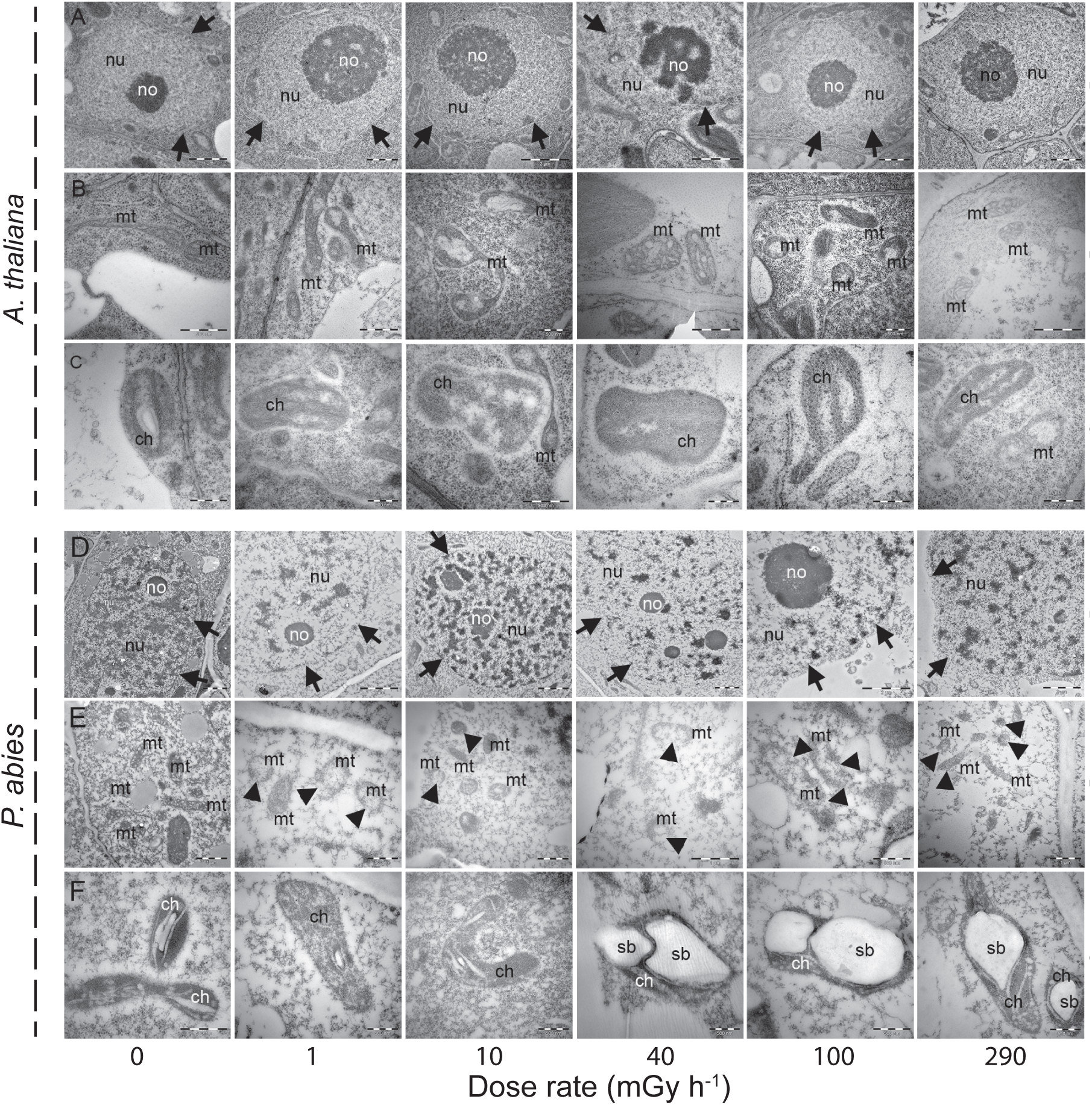
Transmission electron micrographs showing cellular organelles after 48 h of gamma irradiation from day 7 after sowing in (a) – (c) A. *thaliana* and (d) - (e) Norway spruce (*P. abies*). nu: nucleus, no: nucleoli, mt: mitochondrion, ch: chloroplast, sb: starch body. Arrows indicate nuclear membrane. Arrowheads indicate disrupted mitochondria. Five plants were investigated per treatment for each species. Scale bars: Scale nu: 1 µm, ch: 500 nm, mt: 500 nm.

### 3.3 Higher levels of DNA damage in gamma-irradiated Norway spruce than A. thaliana

In *A. thaliana*, the Comet assay showed low but significant damage with ≤ 5% tail DNA at 1-100 mGy h^−1^ and 12% tail DNA at 290 mGy h^−1^ (Fig. 4A). In comparison, DNA damage was consistently 2-8-fold higher in Norway spruce with 10%, 12%, 15%, 17.5% and 40% DNA in the Comet tail after the 48-h of irradiation at 1, 10, 40, 100 and 290 mGy h^−1^, respectively (Fig. 4B). At 77 dpi, the DNA damage was substantially reduced but still observed in *A. thaliana* at ≥ 10 mGy h^−1^ with ≤ 3% tail DNA and in Norway spruce seedlings at ≥ 1 mGy h^−1^, with ≤ 5% tail DNA at all dose rates (Fig. 4A, B).

**Fig. 4.**
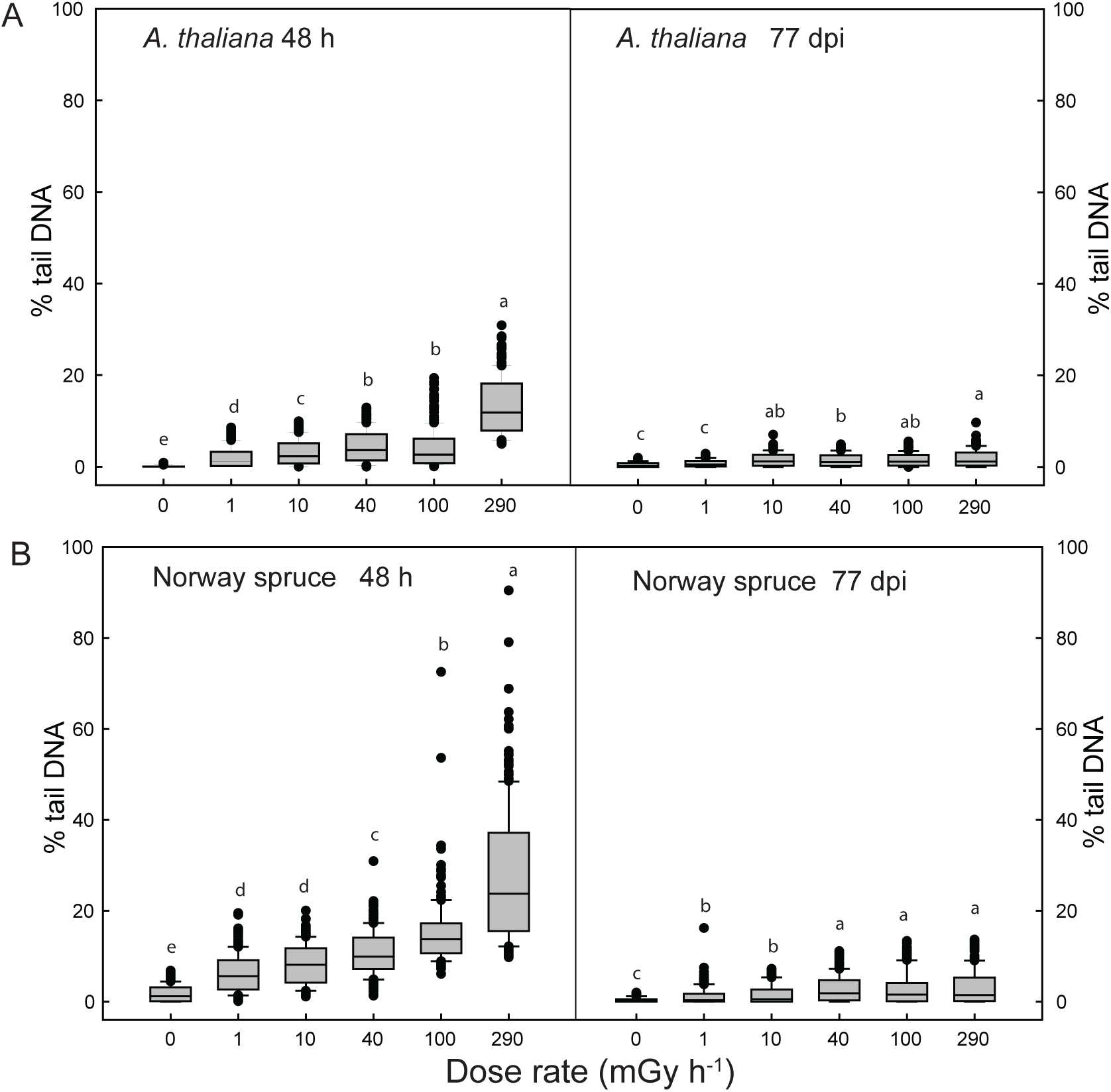
Effect of 48 h of gamma irradiation from day 7 after sowing on DNA damage detected by the COMET assay in (a) *A. thaliana* and (b) Norway spruce after the 48-h irradiation and 77 days post-irradiation (dpi). The line in each box represents the median values of 3 samples with 180-200 nuclei scored in each. Lower and upper box boundaries = 25 and 75 percentiles, error bars = 10 and 90 percentiles with data points outside these shown as dots. Different letters within each figure indicate significant difference (p ≤ 0.05) based on analysis of variance followed by Tukey’s test.

### 3.4. Profound transcriptome differences between gamma-irradiated Norway spruce and A. thaliana

Comparative RNA-seq analysis of gamma-irradiated shoot samples (1-100 mGy h^−1^ both species, 290 mGy h^−1^ *A. thaliana* only), revealed extensive differences between the species. *A. thaliana* expressed a total of 26,643 genes, of which 11,592 were DEGs (43.5%) between any dose rate and the unexposed control in one or multiple gamma dose rates (Fig. 5A, upper panel). Norway spruce expressed a total of predicted 66,069 genes, of which 5326 were DEGs (8.1%, Fig. 5A, lower panel). In *A. thaliana*, a high number of DEGs was observed already from 1 mGy h^−1^ (3442 DEGs), with the higher dose rates having ∼4400-5500 DEGs (Fig. 5A). At 100-290 mGy h^−1^, there was a higher proportion of upregulated genes with more than 2-fold change as compared to the lower dose rates (Fig. 5A). By contrast, in Norway spruce, very few DEGs were observed at 1-10 mGy h^−1^ (38 and 43 DEGs), while 822 and 5047 DEGs were observed at 40-100 mGy h^−1^, respectively (Fig. 5B, C). The increase in DEGs from 40 to 100 mGy h^−1^ was much more pronounced in Norway spruce than in *A. thaliana* (Fig. 5A,). The largest overlap of *A. thaliana* DEGs was between 100 and 290 mGy h^−1^, and between 1, 10 and 40 mGy h^−1^ (Fig. 5B). A good proportion of Norway spruce DEGs at 40 and 100 mGy h^−1^ overlapped, showing some consistency in the response even with a large difference in total numbers of DEGs, including many unique ones (Fig. 5C).

**Fig. 5.**
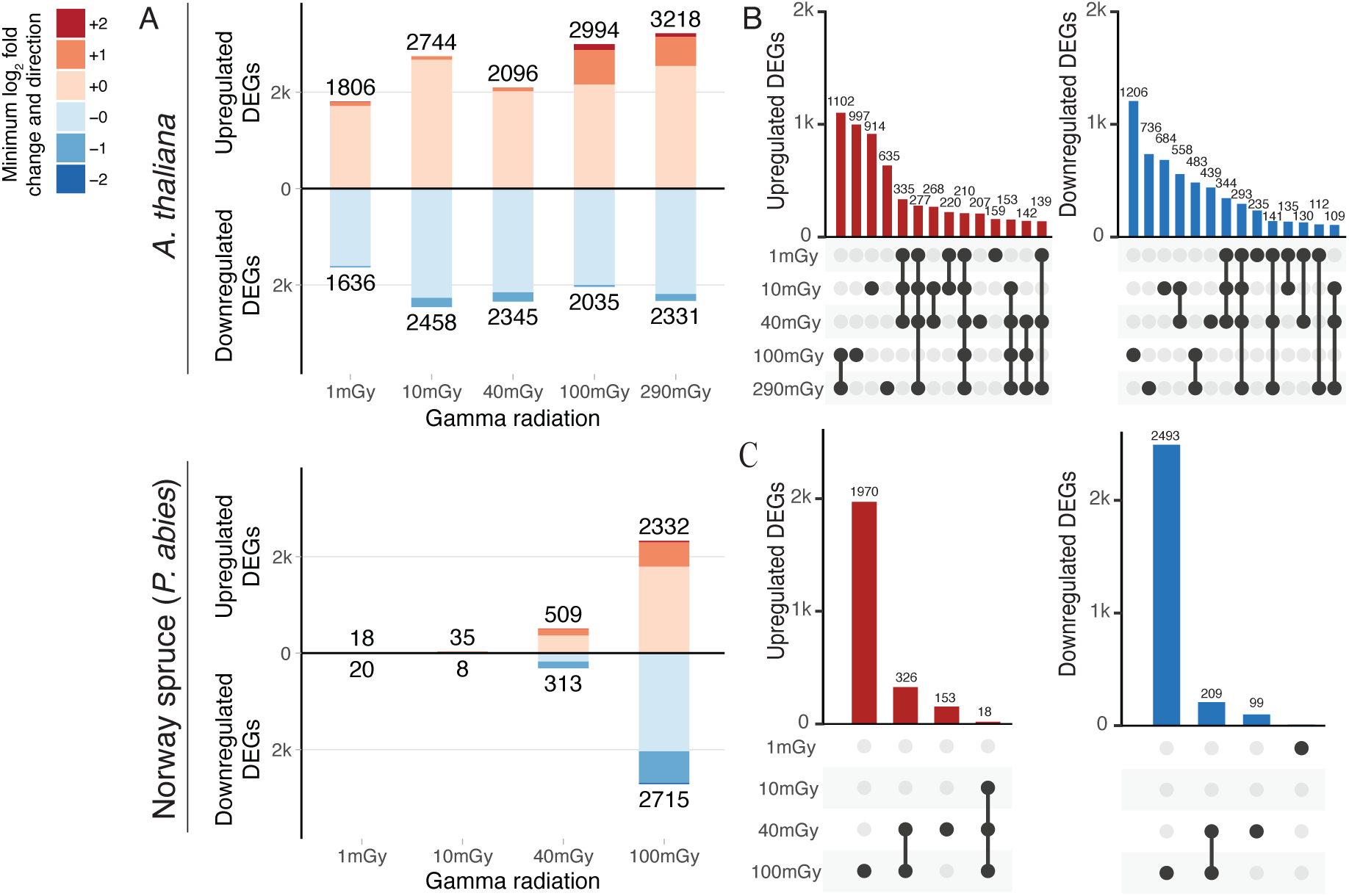
Number of differentially expressed genes (DEGs) and their log_2_ fold change after 48 h of gamma irradiation from day 7 after sowing relative to unexposed controls of (a)-(b). *A. thaliana* and (c)-(d) Norway spruce, with false discovery rate (FDR) < 0.05 (n = 4). Overlap of upregulated (red numbers) and downregulated (blue numbers) genes at different dose rates in (b) *A. thaliana* and (d) Norway spruce. For ea ch species, four repeated samples were analysed in duplicate per treatment.

### 3.5. Distinctive pathway enrichment in Norway spruce and A. thaliana at different dose rates

GO and KEGG pathway analyses on sets of DEGs revealed a gamma dose rate-dependent enrichment of different pathways. In *A. thaliana*, DNA replication, DNA repair, cell cycle, endoreduplication, and endoplasmic reticulum (ER) stress were enriched at 1-40 mGy h^−1^ (Fig. S3, S4). Hormone biosynthesis and signalling pathways showed considerable enrichment along with stress response at 100-290 mGy h^−1^ (Fig. S3, S4). Oxidative stress, defence response, growth and development, protein synthesis and catabolism as well as mitogen-activated protein kinase (MAPK)-mediated signalling were significantly regulated at all dose rates (Fig. S3, S4). In Norway spruce, pathways such as DNA replication, DNA repair, cell cycle, endoreduplication, unfolded protein response (UPR) and ER stress, response to gamma, energy metabolism and growth and development were all enriched at ≥ 40 mGy h^−1^ (Fig. S5, S6). Photosynthesis and protein biosynthesis and metabolism were enriched in downregulated DEGs at 100 mGy h^−1^ (Fig. S5, S6) in Norway spruce only.

In *A. thaliana* KEGG pathway analysis revealed that the DDR pathways such as HR, mismatch repair (MMR), non-homologous end-joining (NHEJ) and base excision repair (BER) were regulated at 10-40 mGy h^−1^, and BER also at 290 mGy h^−1^ (Fig. S4). In Norway spruce, NER, MMR, HR and NHEJ pathways were significantly regulated at 40-100 mGy h^−1^ and BER at 40 mGy h^−1^ only (Fig. S6).

### 3.6. Distinct differences in stress response pathways between Norway spruce and A. thaliana

The specific DEGs in DDR, antioxidant protection and other stress response pathways were further analysed to compare *A. thaliana* and Norway spruce response after the 48-h of gamma irradiation.

#### 3.6.1. DNA damage repair

In *A. thaliana*, most DEGs in the DDR pathways showed a 30-70% change in expression (Table S3). Genes involved in DNA double strand break (DSB) repair via HR, including *RAD51*, *SMC6B* and *BRCA2B*, showed significant upregulation in the 10-290 mGy h^−1^ range. In Norway spruce, most DDR DEGs at 40-100 mGy h^−1^ had at least a 60% change in expression (Table S3), and genes involved in DSB repair via HR; *SRS2*, *RAD52*, *RPA1A*, *MCM8, XRCC2* and *XRCC3*, were all upregulated at 100 mGy h^−1^. A few other genes such as *PARP1* and *PARP2* were significantly upregulated at the highest dose rates in both species. Some DEGs showed opposite changes in expression between species; *MRE11*, *RPA2A/ROR1* and *BRCA1* in the HR pathway, and *MSH7* in the MMR pathway. Unique DEGs to Norway spruce included upregulation of *KU80* at 40-100 mGy h^−1^ and *SOG1* at 100 mGy h^−1^. Unique to *A. thaliana* was downregulation of *ATM* at 100 mGy h^−1^.

#### 3.6.2. Cell cycle and endoreduplication

In *A. thaliana*, most cell cycle-related DEGs, including *CYCBs* and *CDKB*s, were observed at 1-40 mGy h^−1^ but not 100-290 mGy h^−1^ (Table S4). In Norway spruce, no such effects on B type cyclins were observed and most of the cell cycle genes showed changes in expression at ≥ 40 mGy h^−1^. Anaphase-promoting factors *APC6* and *13* and the cyclin-dependent kinases regulatory subunit *CKS1* showed upregulation at ≥ 40 mGy h^−1^. Positive regulators of the cell cycle such as D type cyclins (e.g., *CYCD1*) were significantly upregulated in *A. thaliana* at ≤ 40 mGy h^−1^ but downregulated in Norway spruce at 100 mGy h^−1^. *RBR1* was upregulated in both species but only at 100 mGy h^−1^ in Norway spruce and at most dose rates in *A. thaliana*. *E2FB* was upregulated at 40 mGy h^−1^ in Norway spruce only. *WEE1,* an important G2/M checkpoint regulator, was upregulated at ≤ 40 mGy h^−1^ in *A. thaliana*, but at ≥ 40 mGy h^−1^ in Norway spruce. Most of the cell cycle-related genes, such as *CYC*s and *CDKs* that are also involved in the endocycle, and the endoreduplication-related biological pathway, were enriched in *A. thaliana* at ≤ 40 mGy h^−1^ but fewer such DEGs were observed in Norway spruce (Table S4, S5). A few endoreduplication genes such as *APC5* and *APC6* were upregulated in Norway spruce at 100 mGy h^−1^ but not in *A. thaliana* (Table S5).

#### 3.6.3. Chromatin remodelers and genes in epigenetic pathways

Significant activation of genes involved in the pathway associated with de-condensation of the nucleosome structure and activation of DDR pathways, was observed in Norway spruce at 40-100 mGy h^−1^ (Table S6). By contrast, in *A. thaliana*, several such genes were either downregulated at ≥ 100 mGy h^−1^, or upregulated at ≤ 40 mGy h^−1^ only (Table S6). The DNA helicase *DDM1* was upregulated in *A. thaliana* at ≤ 40 mGy h^−1^ and DNA methyltransferase *CMT3,* histone 2A variant *HTA8* and a component of the core chromatin remodelling complex *ARP6* were significantly downregulated at 100-290 mGy h^−1^ (Table S4, S6). In Norway spruce, histone deacetylase *HOS15* and genes involved in the regulation of DNA methylation, such as *DMT1*, *IDN2* and *SUVH4*, showed significant upregulation at 100 mGy h^−1^.

#### 3.6.4. Hormone biosynthesis and signalling

*A. thaliana* genes involved in biosynthesis and signalling of main growth-promoting hormones auxin, cytokinin, cytokinin, gibberellin (GA) and brassinosteroid, as well as important stress-response-related hormones such as abscisic acid (ABA) and ethylene, were significantly regulated in response to the gamma irradiation (Table S7) especially at higher dose rates. The auxin-responsive MYB transcription factors *MYB93* and *MYB61*, the auxin-responsive genes *IAAs, SAUR10* and *GH3.6*, the auxin biosynthesis gene *YUC9* as well as the auxin efflux carriers and transporters involved in polar auxin transport, *PINs AUX1* and several ABC transporters, were significantly upregulated in this species (Table S7) but downregulated in Norway spruce. Although the auxin-homeostasis-related gene *IAMT1*, and the auxin responsive genes *SAR1* and *SAUR76* showed increased expression at 40-100 mGy h^−1^ in Norway spruce, several auxin signalling genes including *ARF6, TIR1, AIR9* and *IAA26* were significantly downregulated (Table S7). In *A. thaliana*, cytokinin biosynthesis, metabolism as well as cytokinin signalling genes such as *CKX5*, *LOG7, AHP1, SOT8, ARR2* and *ARR3*, were upregulated at 40-290 mGy h^−1^ (Table S7). Contrastingly, none of these genes showed significant regulation in Norway spruce. Similarly, the GA biosynthesis genes *GA2* (*ent-kaurene synthase 1*) and *GA 20-OXIDASE 2* (*GA20OX2),* and the brassinosteroid biosynthesis and homeostasis genes *CYP90C1* and *CYP85A1*, were upregulated in *A. thaliana* but not in Norway spruce.

In general, ABA biosynthesis and signalling genes did not show any gradual up- or downregulation in *A. thaliana* (Table S7). However, the negative regulator of ABA biosynthesis *PUB44*, and the ABA signalling genes *PYL4, CRK29* and *ATHB-6* were downregulated at < 40 mGy h^−1^. Many stress- and ABA-responsive genes such as *ASPG1, PPCK1,* and *RD22* showed consistent increasing expression from 10-290 mGy h^−1^. Among the genes regulated in both species, the ABA signalling gene *CIPK3* showed downregulation in all dose rates in *A. thaliana*, and at 100 mGy h^−1^ in Norway spruce, and *ADH1* were upregulated in both species at higher dose rates. In *A. thaliana*, ethylene biosynthesis and signalling genes were mostly downregulated at lower dose rates with few exceptions (Table S7). Ethylene-responsive genes such as *CTR1, RAP2-2, RAP2-12,* and *ERFs* as well as the ethylene biosynthesis genes *EOL1* and *ACS5* showed significant regulation in *A. thaliana*, but not in Norway spruce.

#### 3.6.5. Genes in antioxidant metabolism pathways

Significant overall upregulation of antioxidant biosynthesis and signalling genes was observed in *A. thaliana* even from 1 mGy h^−1^. By contrast, in Norway spruce, most of the antioxidant genes were not significantly affected below 100 mGy h^−1^ (Table S8). The key flavonoid and anthocyanin biosynthesis genes *CHS* and *DFRA* were upregulated in *A. thaliana* from 1 mGy h^−1^ but significantly up or downregulated in Norway spruce at 100 mGy h^−1^. A flavanol synthase gene (*FLS1*) was upregulated in *A. thaliana* at ≤ 10 mGy h^−1^ and 290 mGy h^−1^ and Norway spruce at 100 mGy h^−1^. The *PER* gene *PER3* was regulated in both species but from lower dose rates in *A. thaliana* than Norway spruce. *GLUTATHIONE-S-TRANFERASEs* (*GSTU/GSTF*) were upregulated in both species.

#### 3.6.6. Cell wall component genes

In *A. thaliana* genes involved in the biosynthesis of cell wall components, cell wall maintenance and fortification, as well as membrane receptors were significantly upregulated at 100-290 mGy h^−1^, commonly with large fold-changes. In Norway spruce, only very few genes related to cell wall biogenesis and maintenance showed regulation, i.e., at 40-100 mGy h^−1^ (Table S9). In *A. thaliana*, *BOR2* involved in cellulose and pectin metabolism, *HHT1* involved in pectin and suberin biosynthesis, the cell wall organisation and biosynthesis genes *FACT* and *CYP86B1*, were upregulated at 100-290 mGy h^−1^. The endoglucanase *GH9C2* involved in cell wall organisation were upregulated in all dose rates. None of these genes showed any significant changes at any dose rate in Norway spruce. A few other genes involved in biogenesis and metabolism of expansins, cellulose and xyloglucan such as the *EXP4* and *8* and *CESA5* were significantly regulated in both species with downregulation in Norway spruce and upregulation in *A. thaliana*, whereas *XTH5* was upregulated in Norway spruce at 100 mG h^−1^

#### 3.6.7. Photosynthesis and energy metabolism

Photosynthesis was massively downregulated in Norway spruce but not in *A. thaliana*. Photosystem (PS) I and II-related genes such as PS I reaction centre subunits *PSAG, PSAK* and *PSAN*, PS I light harvesting complex *LHCA1*, PS II assembly factor *HCF136*, essential enzymes such as NADPH-protochlorophyllide oxidoreductase *PORA*, and Fructose-1,6-bisphosphatase *CYFBP*, chloroplast binding proteins and membrane efflux carriers, were downregulated in Norway spruce at 100 mGy h^−1^ (Table S10). In *A. thaliana,* many genes were upregulated, such as the PS I and II reaction centre components *PSAB* and *PSBB*, and *NDHC* (Table S10).

Similarly, energy metabolism was far more affected in Norway spruce than *A. thaliana* (Table S11). Norway spruce particularly showed upregulation at ≥ 40 mGy h^−1^ of a wide range of genes involved in glycolysis, TCA cycle and gluconeogenesis, such as, Glyceraldehyde-3-phosphate dehydrogenase *GAPC1*, *MALATE DEHYDROGENASE* (*MDH1*), *SUCCINATE DEHYDROGENASE (SDH2-1*), and several homologs of phosphoenolpyruvate carboxykinase *PCKA* and *PFK2*. Additionally, the *CYTOCHROME C1* (*CYTC-1*) and *CYTC OXIDASE 6B1* (*COX6B-1*) genes involved in mitochondrial electron transport were significantly upregulated at 100 mGy h^−1^. *ALCOHOL DEHYDROGENASE 1* (*ADH1*) was significantly upregulated in both species from 100 mGy h^−1^.

### 3.7. Dynamic regulation of repair and protection-related genes in A. thaliana and Norway spruce during gamma irradiation

To evaluate the dynamics of gene expression during radiation stress, both species were exposed to 12, 24 and 48 h of gamma radiation and the transcript levels of selected genes analysed by RT-qPCR (Fig 6, Fig. S7, S8). Consistent with the RNA-seq data, significantly increased transcript levels of the DDR-related *WEE1, PARP2, RAD51, BRCA1* and the cell cycle-related *CYCB1.2* were observed after 48 h in *A. thaliana*, by 5-10-fold as compared to the unexposed controls (Fig. 6, Fig S7). However, after 12 and 24 h exposure no significant effect of the gamma irradiation on their expression was observed. Also, the expression of *SOG1*, *BARD1*, *KU80*, *MSH7*, *PCNA* and *CYCB1.1* did not differ between the controls and any of the gamma dose rates. Of these DDR- and cell cycle-related genes in Norway spruce, the transcript levels of *WEE1* were significantly higher at 20-100 mGy h^−1^ after 24 h only compared to the unexposed controls and those of *SOG1* increased at 40-100 mGy h^−1^ after 24 h, while *PARP2* expression increased significantly by 10-30-fold at 40-100 mGy h^−1^ after 24 and 48 h. There were no significant effects on expression at 12 h.

**Fig. 6.**
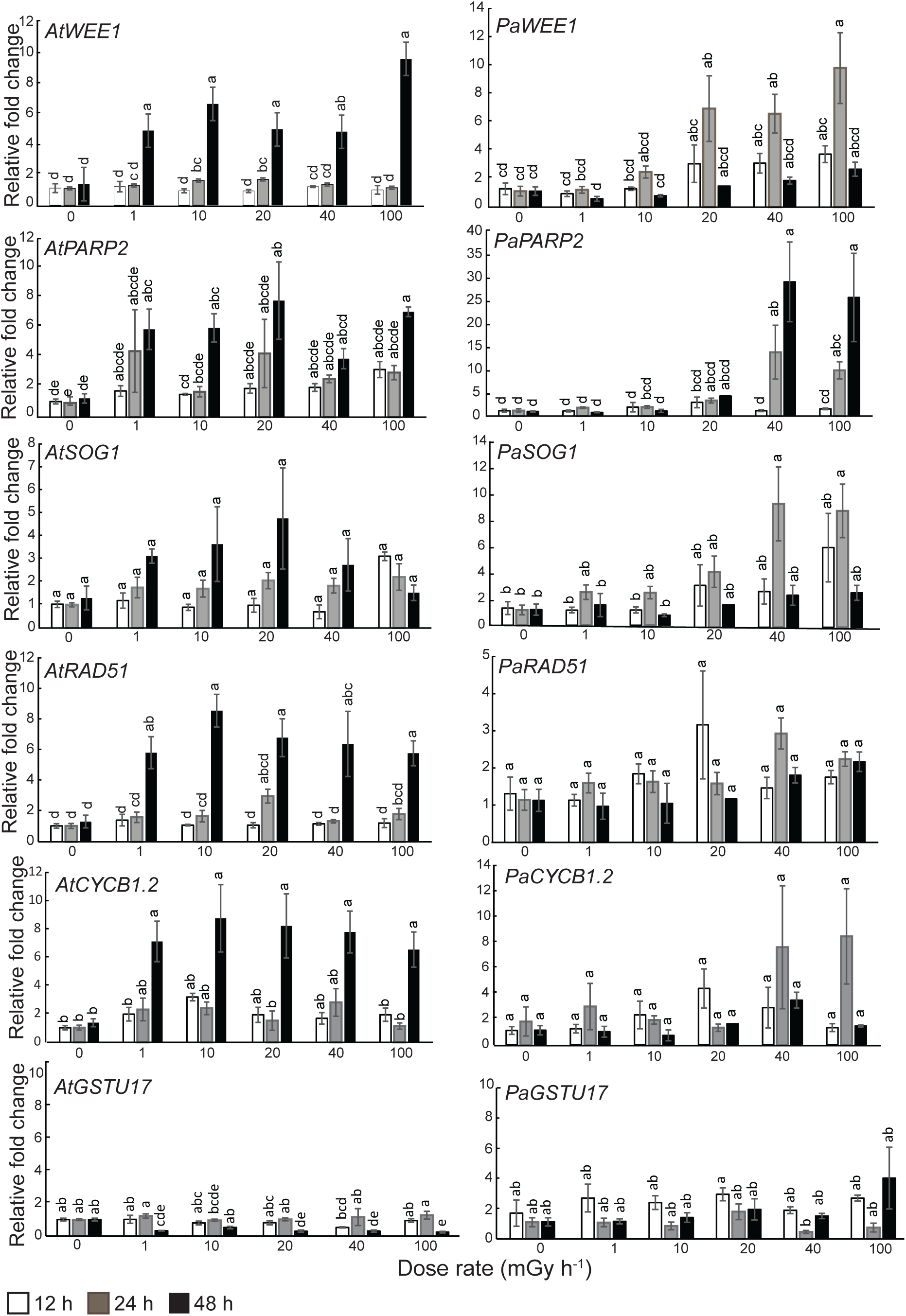
Relative transcript levels of DNA repair and cell cycle genes as well as the antioxidant-related *GLUTATION-S-TRANSFERASE* (*GSTU17*) gene after 12, 24 and 48 h of gamma irradiation from day 7 after sowing in shoots of *A. thaliana (At)* and Norway spruce *(Pa)*. Transcript levels were normalized against *ACTIN*, *ELONGATION FACTOR 1* α and *UBIQUITIN 10* and shown relative to the unexposed controls at 12 h. The results are mean ± SE (n = 3-4 with 3-4 technical replicates). Different letters within each gene indicate significant difference (p ≤ 0.05) based on analysis of variance followed by Tukey’s post-hoc test.

Out of the four antioxidant genes analysed (*DFRA1* and *CHS1* (phenolic/flavonoid biosynthesis)*, GSTU17* and *PER52), GSTU17* showed significant downregulation in *A. thaliana* at 48 h at all dose rates except 10 mGy h^−1^, but no such regulation was observed for Norway spruce (Fig. 6, Fig. S8). Additionally, *CHS1* showed significant upregulation in *A. thaliana* at 12 and 24 h after exposure to 1 and 100 mGy h^−1^ (Fig. S8). Moreover, amongst the two analysed marker genes of retrograde signalling, *AOX1* (alternative oxidase) and *WRKY40* (a transcription factor)*, WRKY40* showed small but significant downregulation and 100 mGy h^−1^ in 12 and 24 h of radiation in *A. thaliana* (Fig. S8). However, no significant regulation of these genes was observed in Norway spruce.

### 3.8. Antioxidants in A. thaliana and Norway spruce during gamma irradiation

To evaluate antioxidant dynamics, the activities/concentrations of CAT, PER, SOD, and flavonoids were investigated in *A. thaliana* and Norway spruce seedlings exposed to 12, 24 and 48 h of gamma radiation (Fig. 7). Although the concentrations of PER and CAT differed significantly between the two species with *A. thaliana* being highest in PER and Norway spruce in CAT, there were generally no significant effects of the gamma irradiation. However, compared to the unexposed controls, the SOD activity in Norway spruce showed significant increase after 48 h at 40-100 mGy h^−1^. There was no effect of gamma radiation after 12 and 24 h radiation in Norway spruce and no significant regulation of SOD in *A. thaliana*. Flavonoid concentrations showed significant downregulation with time in *A. thaliana* regardless of gamma irradiation or not. In Norway spruce, no such effect was observed.

**Fig. 7.**
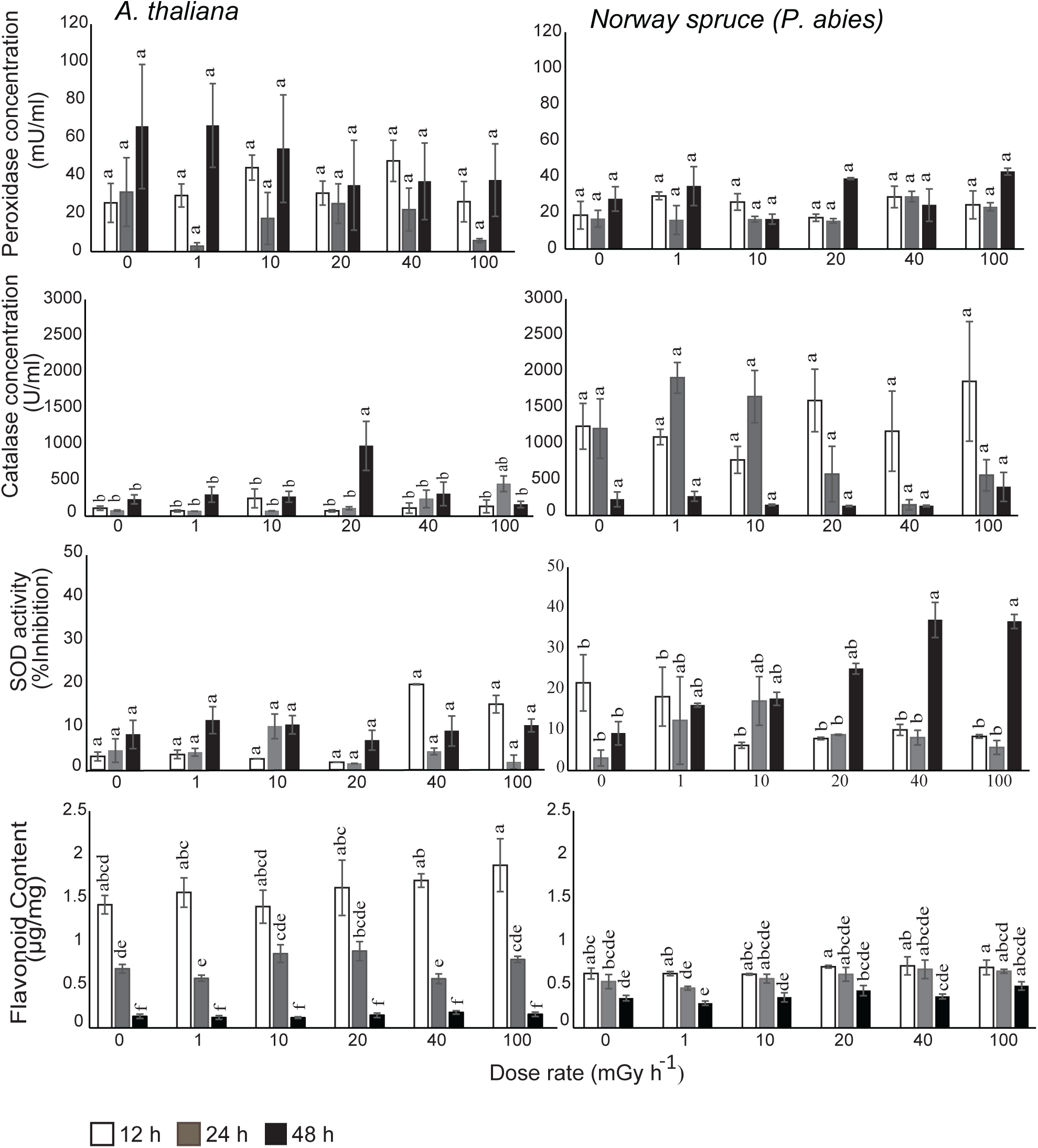
Total antioxidant concentration or activity after 12, 24 and 48 h of gamma irradiation from day 7 after sowing in shoots of *A. thaliana* and Norway spruce (*Picea abies*). The antioxidant results were normalised based on the absorbance of individual standards. The results are mean ± SE (n = 3 or 4 (flavonoids) with 2-3 technical replicates). Different letters within each concentration/activity indicate significant difference (p ≤ 0.05) based on analysis of variance followed by Tukey’s post-hoc test.

## 4. Discussion

To improve the knowledge base for risk assessment and protection of plant communities in radionuclide-contaminated ecosystems against ionising radiation, the current comparative study has focussed on underlying mechanisms with respect to differential radiosensitivity in the highly radiosensitive conifer Norway spruce and the radiotolerant *A. thaliana*. Through exposure of young seedlings to gamma radiation under standardised conditions and comparative assessment of phenotypic, cellular, and genotoxic effects, combined with RNA-seq and studies of dynamics of antioxidant contents and expression of relevant genes, we present novel understanding of differences between the two species in the mechanisms involved in protection and repair.

Previously, we have shown dose rate-dependent impaired development in Norway spruce seedlings exposed to gamma radiation for 144 h, with significant growth inhibition at ≥ 40 mGy h^−1^ and progressively developing after-effects eventually causing mortality (Blagojevic et al. 2019a). By contrast, *A. thaliana* showed no significant adverse effects at dose rates ≤ 540 mGy h^−1^ except a transient delay in reproductive development at the highest dose rates (Blagojevic et al. 2019a). Consistent with the previous findings, in this study *A. thaliana* did not show any clear negative effect on growth after 48 h irradiation up to 290 mGy h^−1^ except a slight transient delay in floral development, whereas Norway spruce showed distinct growth inhibition (Fig. 1, S1). In agreement with this, the RNA-seq analysis revealed profound differences between the two species in response to gamma irradiation (Fig. 5, S3-S6). The fact that *A. thaliana* mounts a transcriptomic response at lower dose rates than Norway spruce signifies that it is more poised towards rapid response to inflicted radiation stress.

The energy deposited at a specific gamma radiation dose rate in the two plant species should be similar, with ionisation and excitation of electrons occurring uniformly throughout the cells. Thus, induced damage should also be randomly distributed in biomolecules such as DNA and relative to the DNA content (genome size) (Einset and Collins 2018). Consistent with a previous report (Blagojevic et al. 2019a) this study showed less DNA damage in *A. thaliana* than in Norway spruce both at the end of the irradiation and post-irradiation period (Fig. 4). We therefore hypothesised that *A. thaliana* has more effective DNA repair and/or protection systems. Yet, the DNA damage at 77 dpi in both species (Fig. 4), albeit at considerably lower level than at the end of the 48-h irradiation, demonstrates that some DNA damage or genomic instability is not critical to survival or development. Although genomic instability may result into detrimental growth effects (Mothersill and Seymour 1998), no clear negative effects on the vegetative growth post-irradiation in *A. thaliana* even after exposure to 400-540 mGy h^−1^ in contrast to the severe growth inhibition and mortality observed in Norway spruce and Scots pine at dose rates ≥ 40 100 mGy h^−1^, (Blagojevic et al. 2019a) confirm better tolerance in *A. thaliana* compared to conifers. In our current study, no mortality after 48 h of gamma irradiation in either species again points to a certain tolerance to damage in both species.

The comparative RNA-seq analysis revealed that both *A. thaliana* and Norway spruce initiated a plethora of DDR pathways to avoid harmful mutations and maintain genome integrity. However, the response of the two species differed in two main aspects (Fig. S9; Table S3). Firstly, the upregulation of DDR-related genes at lower doses dose rates in *A. thaliana* than in Norway spruce suggests that the former is far more responsive towards lower levels of DNA damage. Secondly, the prominent DNA repair pathways in *A. thaliana* were mostly HR, and to some extent MMR, whereas in Norway spruce, in addition to HR, several other pathways such as NHEJ, MMR and DNA transleison synthesis were activated. Hence, it is tempting to speculate that Norway spruce needed to engage a wider range of DDR because of the significantly higher levels of DNA damage compared to *A. thaliana*, possibly because of less antioxidant protection and a larger genome.

One of the key contributor genes in gamma radiation-induced DDR in plants*, SOG1*, which activates several DDR pathways and cell cycle checkpoints (Kim et al. 2019), was not upregulated at any time points or dose rates in *A. thaliana* (Fig. 6, Table S3). This might indicate that induction of *SOG1* expression is only modulated in higher total doses or acute radiation that generate very high level of DNA damage (Bourbousse et al. 2018). However, no change in *SOG1* expression after 48 h of gamma irradiation does not necessarily indicate the lack of SOG1 activity as that could rather depend on the post-translational modification by ATM- or ATR-kinases (Yoshiyama et al. 2013). The involvement of *SOG1* in *A. thaliana* is indeed supported by the activation of many primary target genes of SOG1 (i.e., *RAD51, BRCA1, PARP1* and *2, GMI1*) in response to the 48 h of gamma irradiation. By contrast, significant upregulation of *SOG1* in Norway spruce after 24 h at ≥ 40 mGy h^−1^ and 48 h at 100 mGy h^−1^ (Fig. 7, Table S3) but no significant regulation of *ATM* (Table S3), might imply that elevated *SOG1* expression was triggered in Norway spruce possibly due to higher levels of accumulated DNA damage. Moreover, upregulation of *WEE1,* a direct target of SOG1 and key regulator of CDK-inhibition and stress-induced cell cycle arrest (Kevei et al. 2011; De Schutter et al. 2007) (Fig. 6, Table S4), coincided with *SOG1* expression in Norway spruce, strongly suggesting SOG1 involvement in gamma-induced DDR and cell cycle arrest (Pedroza-Garcia et al. 2022; Bourbousse et al. 2018).

Transcriptional activation of numerous B-type *CYC*- and *CDKB*-genes in *A. thaliana* after 48 h at 1-40 mGy h^−1^ (Fig. 6, Table S4) might indicate gamma radiation stress-induced cell cycle arrest and subsequent DDR through HR by maintaining higher cyclin threshold (Endo et al. 2006). This further supports the previous notion that enhanced CDKB1 activity is involved in ionising radiation-induced cell cycle arrest leading to HR-repair in damaged mitotically active cells (Weimer et al. 2016). On the contrary, there were fewer regulations and downregulation only of B-type *CYC* or *CDK* genes in Norway spruce (Table S4), indicating no significant involvement of such genes in such cell cycle arrest. The regulation of the key contributors in cell cycle progression in both species, albeit to a larger degree in *A. thaliana*, suggests that the cell cycle is one of the most affected biological pathways in the applied gamma radiation. Most of the CYCs and CDKs regulated in *A. thaliana* in our study are also involved in the regulation of the endocycle. However, stress-indued endoreduplication is less common in gymnosperms than angiosperms (De Veylder et al. 2001) and fewer genes related to the endocycle were upregulated in Norway spruce compared to *A. thaliana* (Table S4, S5). Increased endoreduplication was previously shown in *A. thaliana* at 1 Gy min^−1^ (150 Gy totally) and *Lemna minor* at 1500 mGy h^−1^ for 7 days, but not at lower dose rates (Adachi et al. 2009; Van Hoeck et al. 2015). Our preliminary study did not reveal any effect of 48 h gamma radiation at 100 mGy h^−1^ on endoreduplication in *A. thaliana* and Norway spruce (flow cytometry results not shown), suggesting that the applied dose was not sufficient to induce the endocycle.

DDR is closely related to epigenetic regulation and chromatin structure must be decondensed for the DDR proteins to carry out efficient repair (Choi et al. 2019; Kim et al. 2019). Chromatin remodelers were previously shown to be upregulated in *A. thaliana* at very high dose rates of ionising radiation (X-rays at 5.25 Gy h^−1^ with total doses of 10-100 Gy) (Sidler et al. 2015). However, in our study upregulation of such genes in *A. thaliana* occurred at ≤ 40 mGy h^−1^ only. By contrast, the upregulation of *H2A* variants, DNA methylation and deacetylation-related genes in Norway spruce at ≥ 40 mGy h^−1^ only (Table S5, Fig. S5) clearly indicates chromatin modification under high radiation stress in this species. The enrichment of DNA methylation regulation is consistent with the previously reported DNA hypermethylation in *P. sylvestris* trees exposed to high doses of ionising radiation in the Chernobyl area, a mechanism suggested to stabilise the genome (Kovalchuk et al. 2003). Despite the genome of Norway spruce being 150 times bigger than that of *A. thaliana*, it only contains 2.5 times more predicted genes (Nystedt et al. 2013), and has large heterochromatin regions which requires the activation of chromatin remodelers for decondensation for efficient DDR (Fig. S9). Moreover, gamma radiation-induced damage to heterochromatin regions does not initiate rapid activation of DDR pathways (Falk et al. 2008). It is thus plausible that Norway spruce requires significantly more DNA damage for the activation of DDR and chromosome remodelers.

The 48-h exposure did not affect the structural integrity of the SAMs (Fig. 2) in contrast to the previous studies where 144 or 360-h gamma irradiation resulted in adverse effects in SAMs of Norway spruce only (Blagojevic et al. 2019a). However, TEM image analysis revealed that integrity of thylakoid and nuclear membranes was compromised in Norway spruce, but not in *A. thaliana* (Fig. 3). Notably, in both species the mitochondria appeared to be the most vulnerable organelle. Furthermore, the fact that disrupted mitochondria were evident from 1 mGy h^−1^ in Norway spruce, while *A. thaliana* showed no damage below 100 mGy h^−1^, indicates a major difference in their capacity to maintain cell homeostasis.

Former studies have demonstrated activation of ROS-scavenging enzymes such as SOD and APX in gamma-irradiated plant species such as *A. thaliana* and *L. minor* (Vanhoudt et al. 2014; Xie et al. 2019) and flavonoids in *Onobrychis viciifolia* and *Glycine max* (Katiyar et al. 2022). Lack of activation of antioxidant-related genes in Norway spruce at ≤ 40 mGy h^−1^ but comprehensive upregulation of such genes (*PERs, CHSs, GSTUs*, anthocyanin and flavanol synthesis genes) at all dose rates in *A. thaliana* (Table S8), may at least partly explain the damage to mitochondria, chloroplasts, and nuclear membranes at all dose rates in Norway spruce in contrast to only slight mitochondrial damage at high dose rates in *A. thaliana* (Fig. 3). Despite the observed organelle damage, the lack of significant upregulation of two marker genes for retrograde signalling, *AOX1* and the transcription factor *WRKY40,* at all dose rates in both species (Fig S8), may possibly suggest that the duration or levels of the gamma irradiation were insufficient to activate retrograde signalling.

In the time-kinetic analyses of antioxidants (PER, CAT, flavonoids, SOD) in this study, only SOD in Norway spruce exhibited increased activity in response to gamma irradiation (after 48 h at 40 and 100 mGy h^−1^; Fig. 7). This suggests that the dynamics of the formation of these antioxidants in response to radiation stress may vary among different species. Additionally, increased biosynthesis of antioxidants may require a longer period of gamma irradiation. At least the lack of increase in flavonoids in Norway spruce and *A. thaliana* in this study (Fig. 7) and in Scots pine after 144 h of gamma irradiation (Blagojevic et al. 2019b) but elevated flavonoid levels in Scots pine under chronic high radiation in the Chernobyl area (Nybakken et al. 2023) are consistent with requirement of long-term irradiation. It is also possible that assays of total antioxidant content/activity might not detect nuances of antioxidant homeostasis provided by individual members of the antioxidant families, including the antioxidant status in different cellular compartments.

Moreover, antioxidants such as PERs can modulate cell wall composition during stress (Díaz-Tielas et al. 2012). The cell wall components cellulose and pectin were previously shown to be damaged by very high doses of gamma radiation (1-2.5 kGy) (Kovács and Keresztes 2002). Hence, in contrast to Norway spruce, the radical upregulation of cell wall component genes even at lower gamma dose rates in *A. thaliana* (Table S9) might suggest efficient replacement of damaged cell wall components and cell wall fortification in *A. thaliana* (Le Gall et al. 2015).

Antioxidants such as *GST*s may also downregulate auxin-responsive genes (Sylvestre-Gonon et al. 2019; Jiang et al. 2010). The upregulation of *GSTs* and growth retardation in Norway spruce at higher dose rates as well as downregulation or no regulation of growth-promoting hormone pathways are consistent with this notion (Fig.1, S1, Table S7). By contrast, the gamma-irradiation-induced upregulation of auxin and cytokinin-related genes (biosynthesis, metabolism, signalling, auxin transport) in *A. thaliana* complies with the maintenance of meristem function to sustain normal growth. These results are in line with the previously demonstrated activation of such genes in response to abiotic stress, indicating the importance of growth-hormone homeostasis under stress (Verma et al. 2016). Similar to previous reports on the induction of ABA biosynthesis and signalling genes triggered by ROS in gamma-irradiated *A. thaliana* (Vishwakarma et al. 2017), upregulation of many such ABA-responsive genes were observed in *A. thaliana* in our study, although the total doses applied were much lower. Also, upregulation of the ethylene-responsive oxidative stress-pathway (*ERFs*, *ACS5*) and SA-responsive genes (*MYBs*) in *A. thaliana*, but not Norway spruce (Table S7), indicate less induction of hormone response to gamma-stress in the latter and is consistent with the greater tolerance of *A. thaliana* to gamma radiation. Hence, in *A. thaliana* better antioxidant homeostasis, cell wall biosynthesis/fortification and hormone signalling might contribute to the high tolerance to the gamma irradiation. Conversely, less transcriptional activation of the same protective pathways in Norway spruce possibly have resulted in adverse effects on overall growth and higher susceptibility to radiation-induced oxidative damage.

The RNA-seq analysis provided evidence that energy metabolism and photosynthesis in Norway spruce were strongly affected by gamma irradiation in Norway spruce, but only marginally in *A. thaliana*. In Norway spruce, the reduced expression of photosynthesis-related genes coincided with the reduced growth and chloroplast impairment at higher dose rates (≥ 40 mGy h^−1^) (Table S10). Concurringly, the strong upregulation of energy metabolism-related genes (glycolysis, Krebs cycle, mitochondrial electron transport-related genes) (Table S11) in this species may support the hypothesis that there is a high ATP requirement for damage repair including maintenance of the large genome. Hence, it appears that radiation induces energy deficiency, which in turn requires increased energy expenditure on damage repair at the expense of growth and is reflected by the downregulation of growth-promoting hormones in Norway spruce at higher dose rates. However, the upregulation of energy metabolism genes might additionally indicate that the damage to mitochondria at high dose rates could have led to inefficient energy production and thus a need for the activation of respiratory genes to meet the energy demands.

## 5. Conclusions

This study demonstrates that compared to the radiosensitive Norway spruce, the radiotolerant *A. thaliana* mobilises more comprehensive gamma radiation-induced defence, repair, and homeostasis-maintaining stress responses, even at low dose rates given for 48 h. Despite significant adverse effects in Norway spruce related to DNA and organelle damage and growth at low dose rates, transcriptional activation of DDR was evident at 40-100 mGy h^−1^ only. Furthermore, in Norway spruce, mobilisation of antioxidant genes, as well as modulation of growth-regulatory and stress-related hormones at lower dose rates were less efficient than in *A. thaliana*. Thus, the short-lived *A. thaliana* with short generation-time, and small genome with less energy requirement for maintenance, shows better plasticity in response to radiation stress compared to the long-lived Norway spruce with a lengthy generation time and gigantic genome.

It should be noted that, although sequence similarities indicate similar gene functions of specific genes in Norway spruce and *A. thaliana*, very few genes in Norway spruce have been functionally validated. This is largely due to constraints such as the giant, hitherto relatively poorly characterised genome as well as the lack of efficient genetic transformation and mutagenesis methods for this species. Furthermore, given the substantial variation in radiosensitivity across plant taxa (Duarte et al. 2023), comparative analyses involving a broad range of species are necessary to gain a more comprehensive understanding of the mechanisms underlying differential radiosensitivity. Nevertheless, the observations in this study expand our understanding of differential radiosensitivity in plants and enhance the knowledge base for risk assessment and protection of ecosystems against ionising radiation. In the context of environmental impact, the findings confirm that under severe ecosystem exposure, radiosensitive conifers require special attention with respect to remediating measures.

## Supporting information

Figure legends for supplementary figures

Table legends for supplemental data

## Acknowledgments

This study has been funded by the Research Council of Norway through its Centre of Excellence (CoE) funding scheme (Project No. 223268/F50) and the Norwegian University of Life Science (NMBU). Sincere thanks to Marit Siira and Svein Andre Kolltveit for skilful assistance in plant growing and growth measurements and Tone Ingeborg Melby and Linda Ripel for skilful work with qPCR analyses of gene expression.

## Author contributions

Payel Bhattacharjee: Conceptualization, Data curation, Formal analysis, Investigation, Methodology, Software, Validation, Visualization, Writing - original draft, Writing - review & editing; Dajana Blagojevic: Conceptualization, Data curation, Formal analysis, Investigation, Methodology, Writing – review and editing; YeonKyeong Lee: Methodology, Visualization, Formal analysis, Writing – review and editing; Gareth B Gillard: Data curation, Formal analysis, Visualization, Writing - review & editing; Lars Grønvold, Torgeir R Hvidsten, Simen R Sandve: Data curation, Formal analysis, Writing – review and editing; Ole C Lind: Conceptualization, Writing – review and editing; Brit Salbu : Resources; Funding acquisition, Writing – review and editing; Dag A Brede: Investigation, Methodology, Writing - review & editing, Jorunn E Olsen: Conceptualization, Funding acquisition, Investigation, Methodology, Project administration, Resources, Supervision, Validation, Visualization, Writing - original draft, Writing - review & editing.

## Data availability

The raw RNA sequence data are available in ArrayExpress under the accession number E-MTAB-8081 for Norway spruce and E-MTAB-11120 for *A. thaliana.* Other data that support the findings of this study are available within the article, in the Supporting Information of this article and in online repositories (https://gitlab.com/garethgillard/CompRad-Gamma). The names of the online repository and the link of the identifiers will be available in the Norwegian University of Life Sciences archive once the submission is approved.

