## Supplementary figures and images for "High radiosensitivity in the conifer Norway spruce (*Picea abies*) due to less comprehensive mobilisation of protection and repair responses compared to the radiotolerant *Arabidopsis thaliana*"

### Fig. S1

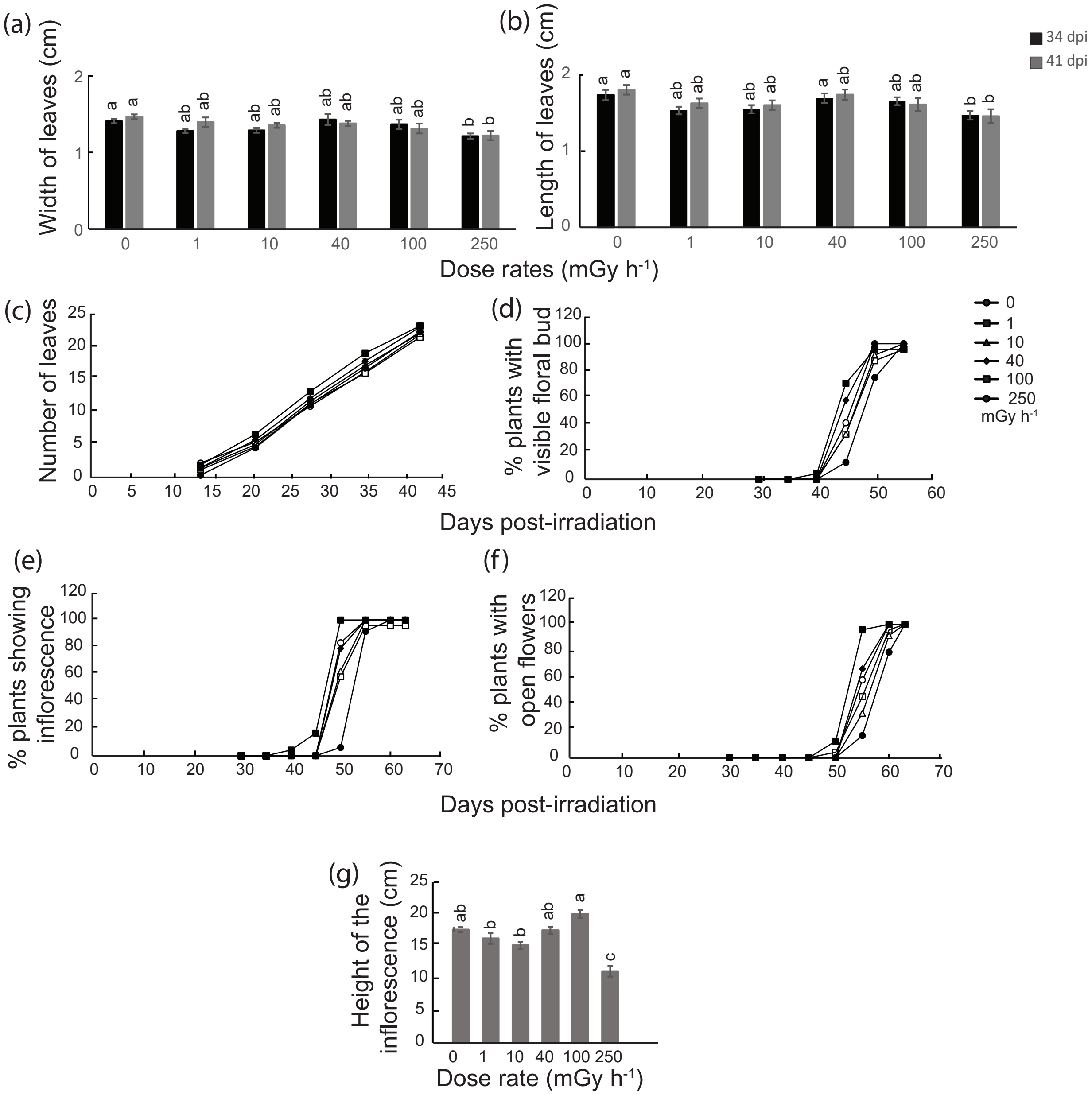


Supplemetary Fig. S1 *Bhattacharjee et al. 2025*

### Fig. S2

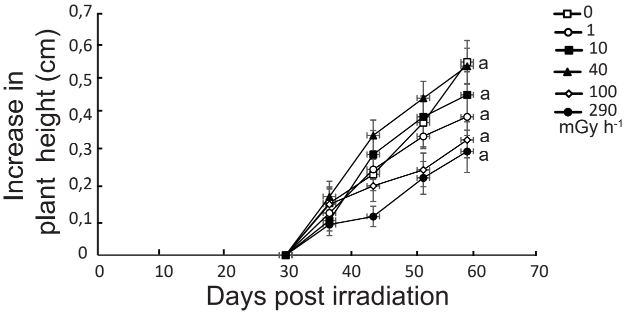


Supplemetary Fig. S2 *Bhattacharjee et al. 2025*

### Fig. S3

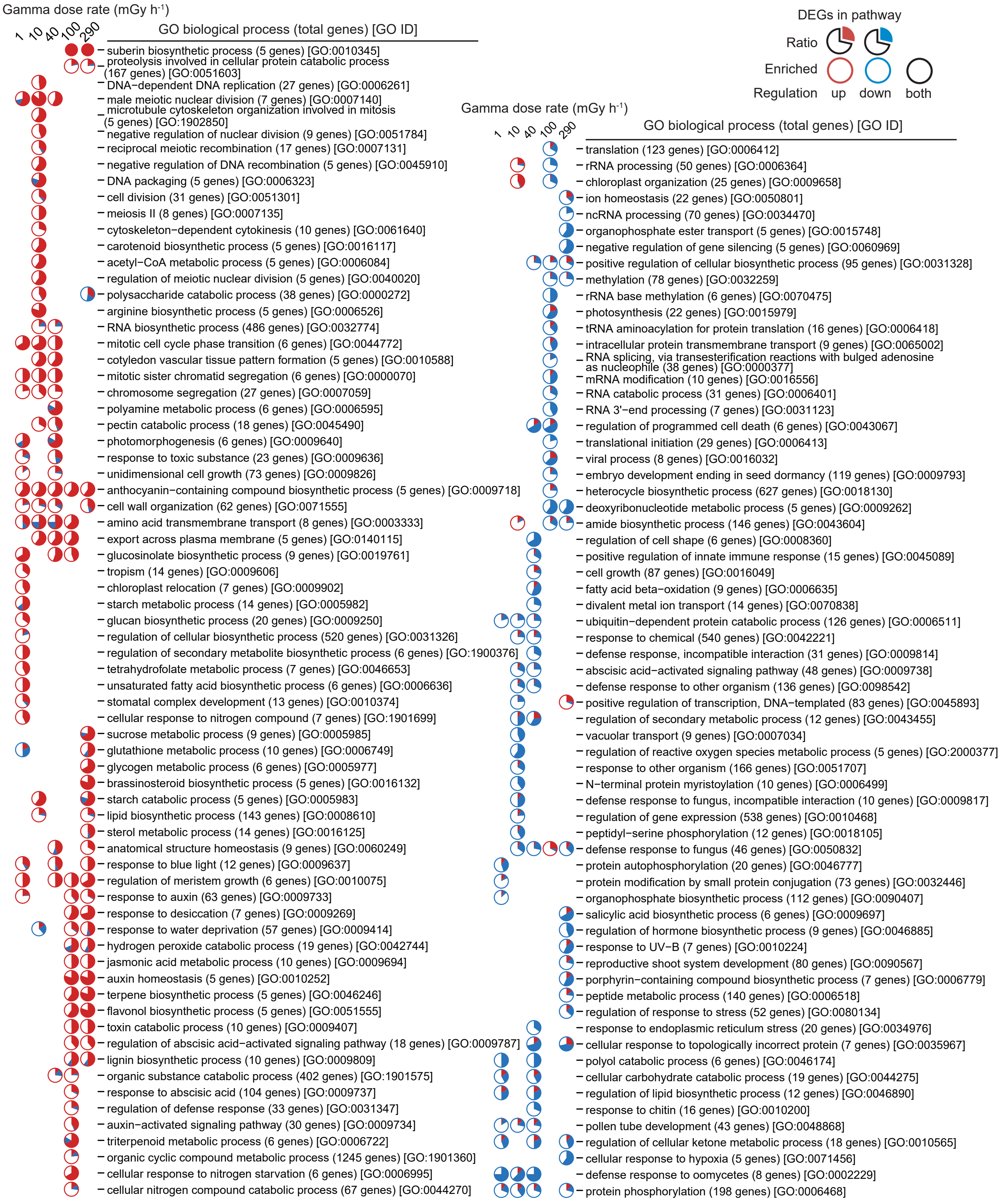


Supplemetary Fig. S3 *Bhattacharjee et al. 2025*

### Fig. S4

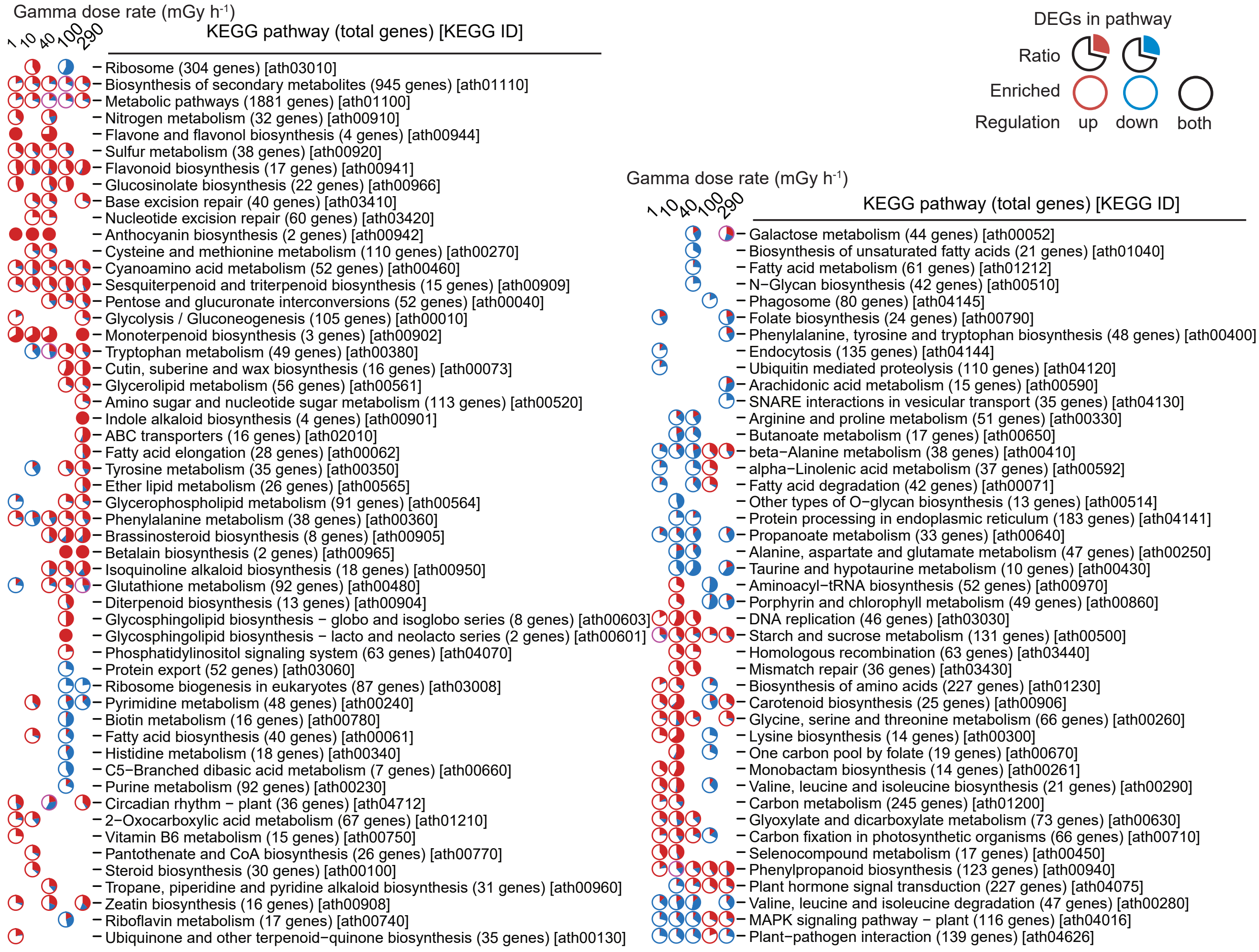


Supplemetary Fig. S4 *Bhattacharjee et al. 2025*

### Fig. S5

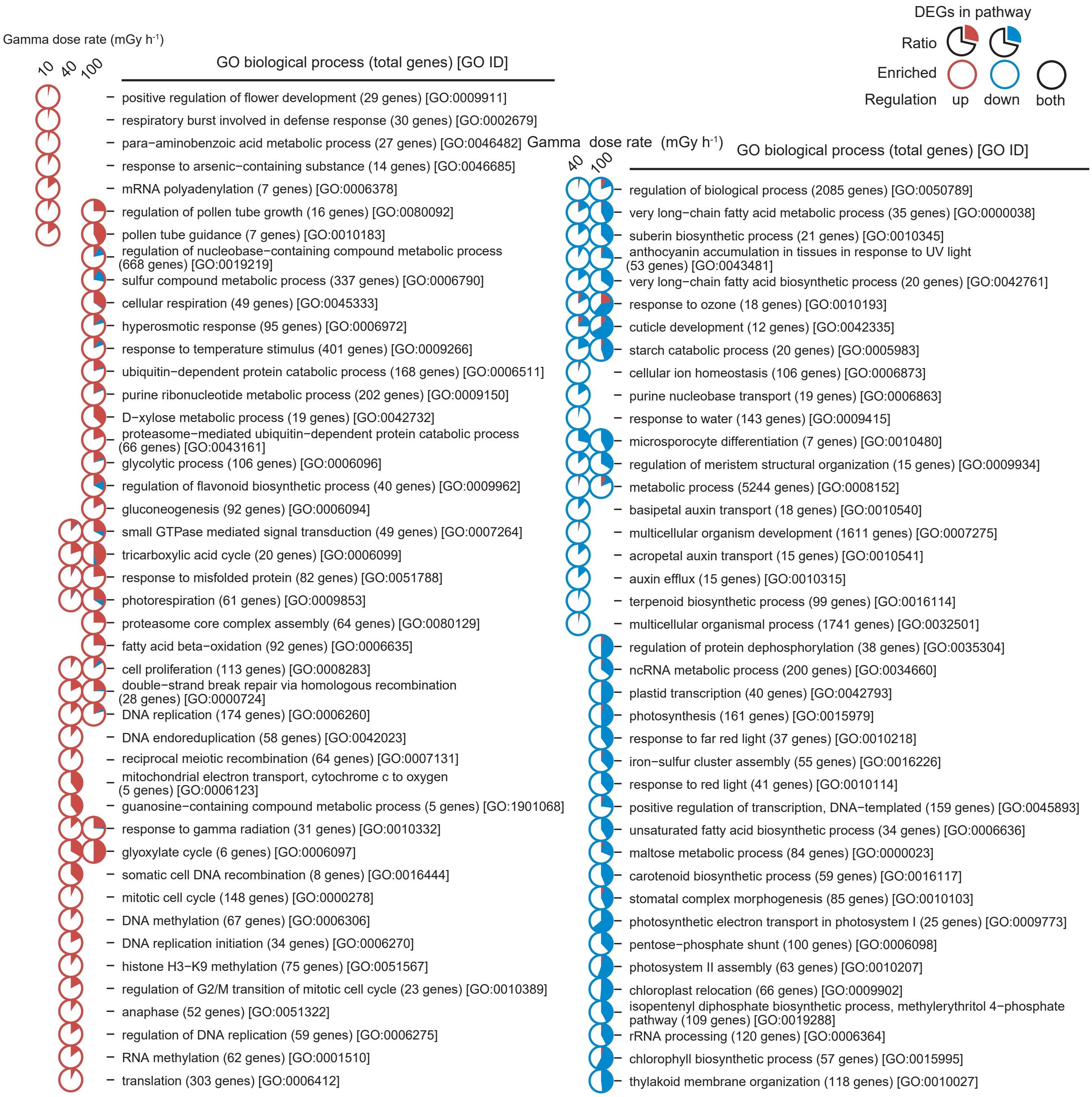


Supplemetary Fig. S5 *Bhattacharjee et al. 2025*

### Fig. S6

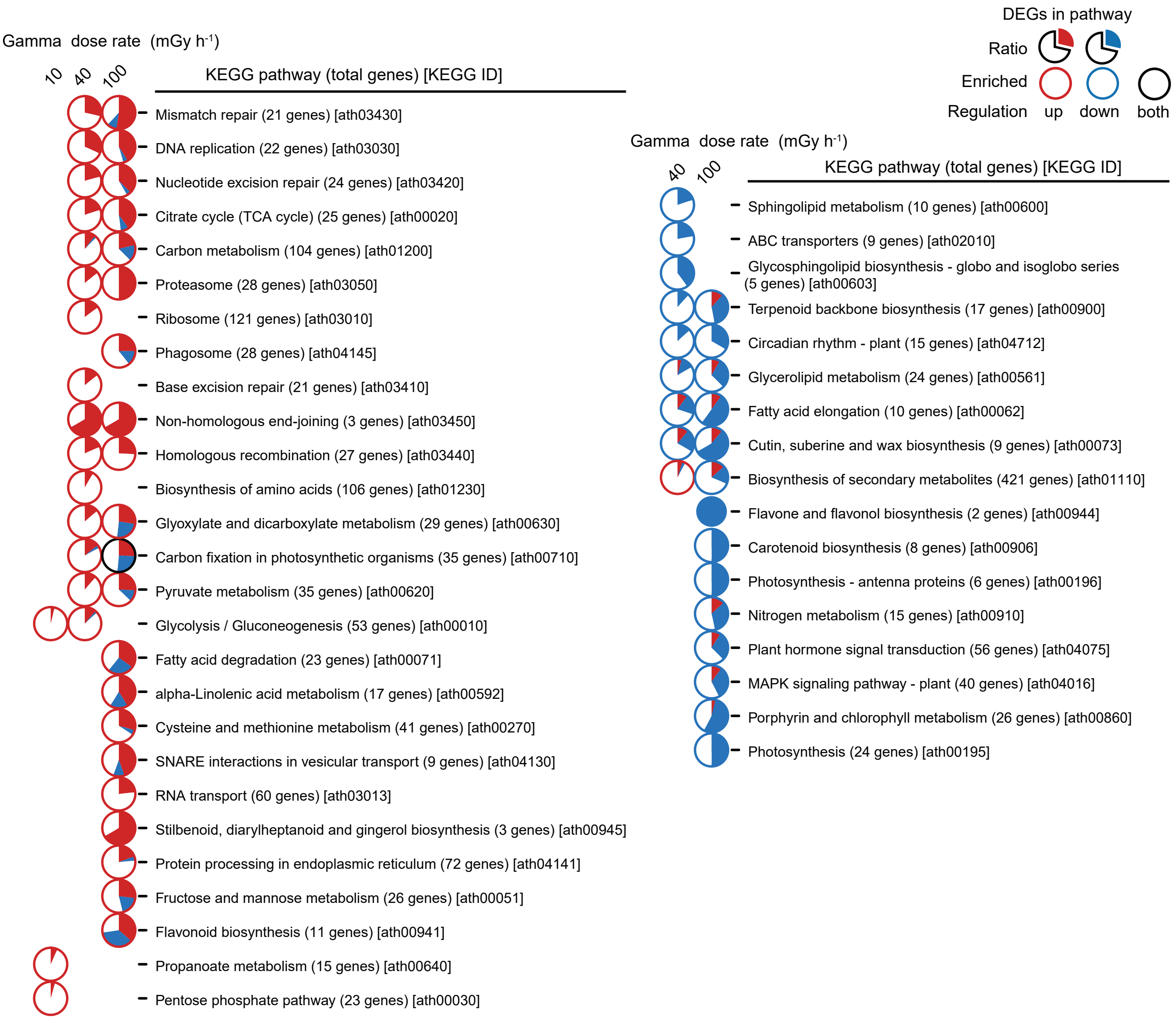


Supplemetary Fig. S6 *Bhattacharjee et al. 2025*

### Fig. S7

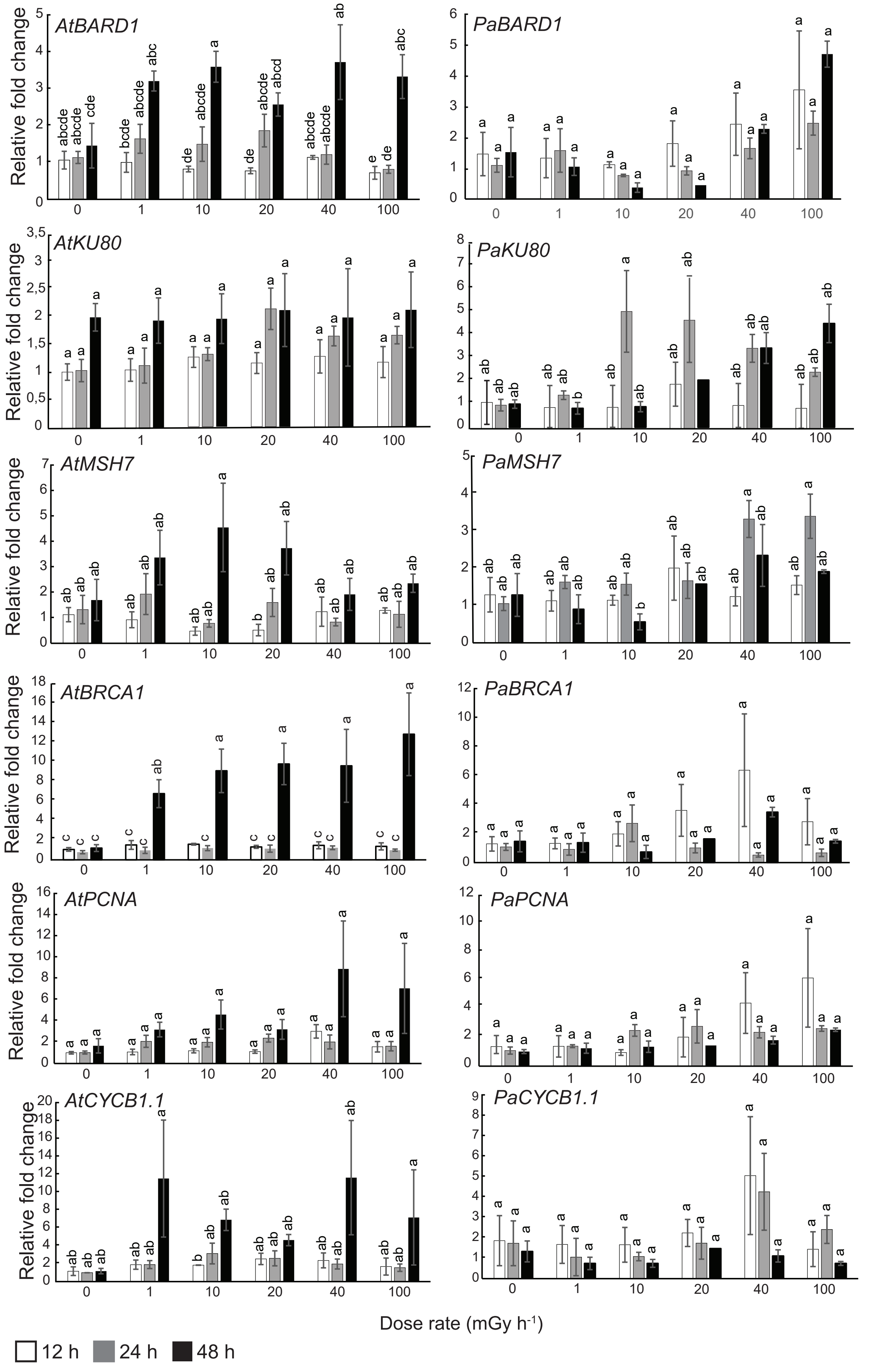


*Fig. S7  Bhattacharjee et al. 2025*

### Fig. S8

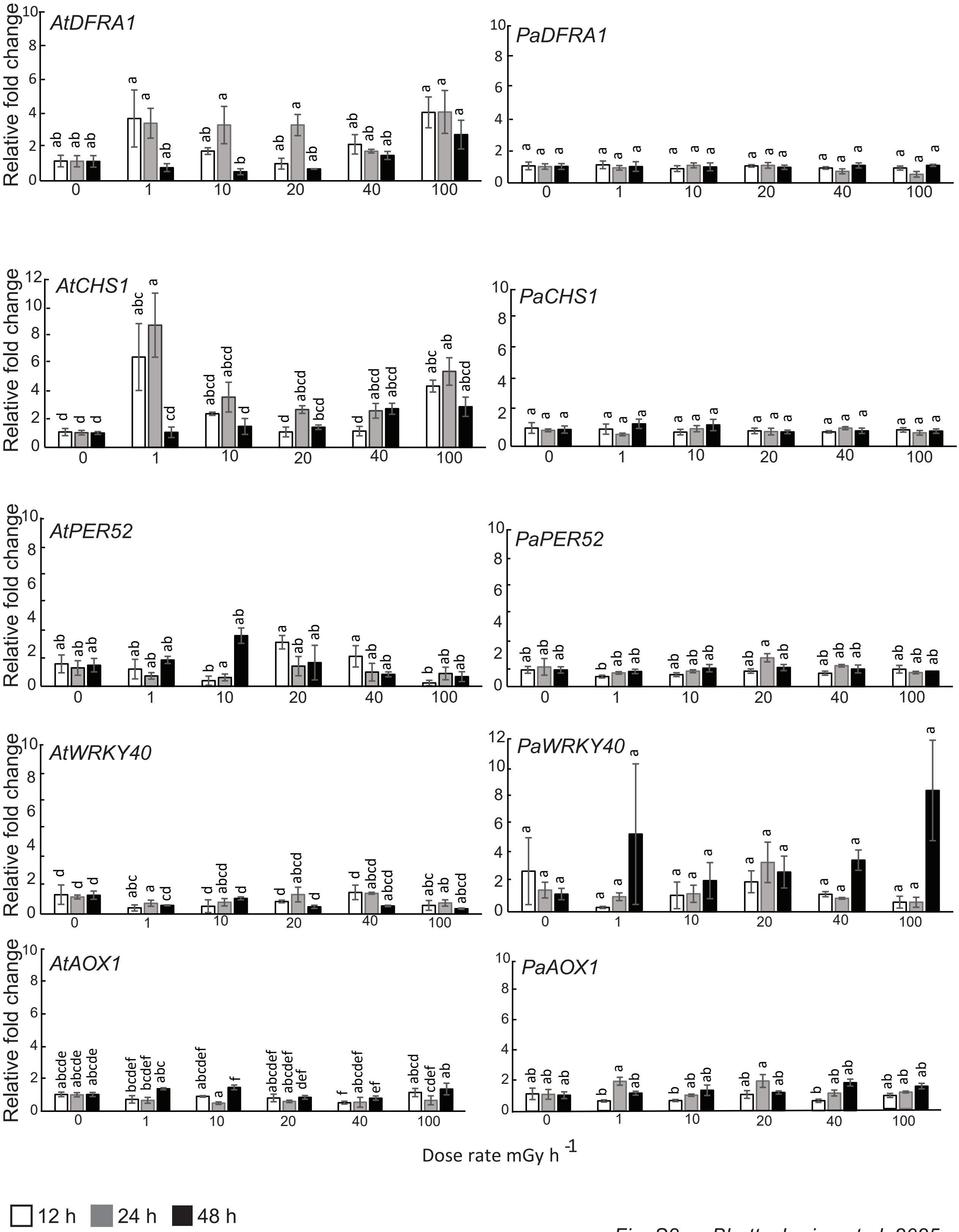


*Fig. S8 Bhattacharjee et al. 2025*

### Fig. S9

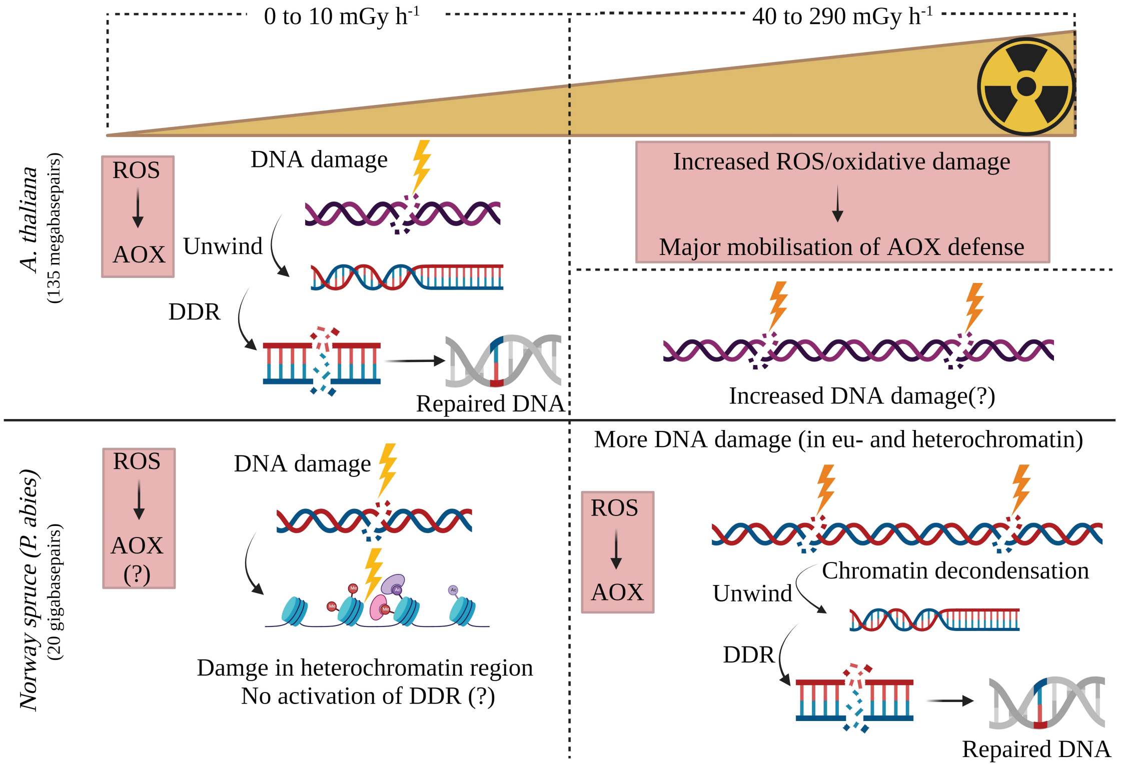


Fig. S9 *Bhattacharjee et al. 2025*
