## Supplementary material for "High radiosensitivity in the conifer Norway spruce (*Picea abies*) due to less comprehensive mobilisation of protection and repair responses compared to the radiotolerant *Arabidopsis thaliana*": Figure legends for supplementary figures

**Fig. S1.** Post-irradiation growth and development of *A. thaliana*. (a) Leaf length and (b) leaf width 34 and 41 days post-irradiation (dpi), (c) number of rosette leaves in a time course, (d-f) different parameters of floral development up to 69 dpi and (g) height of inflorescence 64 dpi (n = 23-24 plants). Means ± standard errors are shown. Different letters within a figure denotes significant difference (p ≤ 0.05) based on analysis of variance followed by Tukey`s test.

**Fig. S2.** Cumulative growth of Norway spruce post irradiation (n = 11-22). Means ± standard errors are shown.

**Fig. S3.** Gene ontology (GO) enrichment analysis of the differentially expressed genes (DEGs) in *A. thaliana* after 48 h of gamma irradiation from day 7 after sowing. The circles represent enrichment with filled colours showing the proportion of upregulated (red) or downregulated (blue) genes out of the total enriched DEGs in a pathway. Per treatment, four repeated samples were analysed in duplicate.

**Fig. S4.** Kyoto Encyclopedia of Genes and Genomes (KEGG) pathway enrichment analysis of the differentially expressed genes (DEGs) in *A. thaliana* after 48 h of gamma irradiation from day 7 after sowing. The circles represent enrichment with filled colours showing the proportion of upregulated (red) or downregulated (blue) genes out of the total enriched DEGs in a pathway. Per treatment, four repeated samples were analysed in duplicate.

**Fig. S5.** Gene ontology (GO) enrichment analysis of the differentially expressed genes (DEGs) in Norway spruce after 48 h of gamma irradiation from day 7 after sowing. The circles represent enrichment with filled colours showing the proportion of upregulated (red) or downregulated (blue) genes out of the total enriched DEGs in a pathway. Per treatment, four repeated samples were analysed in duplicate.

**Fig. S6.** Kyoto Encyclopedia of Genes and Genomes (KEGG) pathway enrichment analysis of the differentially expressed genes (DEGs) in Norway spruce after 48 h of gamma irradiation from day 7 after sowing. The circles represent enrichment with filled colours showing the proportion of upregulated (red) or downregulated (blue) genes out of the total enriched DEGs in a pathway. Per treatment, four repeated samples were analysed in duplicate.

**Fig. S7.** Relative transcript levels of DNA repair and cell cycle genes in *A. thaliana* and Norway spruce after 12, 24 and 48 hours of gamma irradiation from day 7 after sowing. Transcript levels were normalized against *ACTIN*, *ELONGATION FACTOR 1 α* and *UBIQUITIN 10* and shown relative to the unexposed controls at 12 h. The results are mean ± SE (n = 3-4 with 3-4 technical replicates). Different letters within each gene indicate significant difference (p ≤ 0.05) based on analysis of variance followed by Tukey’s post-hoc test.

**Fig. S8.** Relative transcript levels of antioxidants and retrograde signaling marker genes in *A. thaliana* and Norway spruce after 12, 24 and 48 hours of gamma irradiation from day 7 after sowing. Transcript levels were normalized against *ACTIN*, *ELONGATION FACTOR 1 α* and *UBIQUITIN 10* and shown relative to the unexposed controls at 12 h. The results are mean ± SE (n = 3-4 with 3-4 technical replicates). Different letters within each gene indicate significant difference (p ≤ 0.05) based on analysis of variance followed by Tukey’s post-hoc test.

**Fig. S9**. A graphical summary of the plausible molecular effects of gamma radiation exposure on *A. thaliana* and Norway spruce *(P. abies*). The left panel describes the effects of low dose rates (0 to 10 mGy h^-1^) and the right panel describes the effect of higher dose rates (40 to 290 mGy h^-1^) in both species. ROS: reactive oxygen species, AOX: antioxidants, DDR: DNA damage repair.
