## Supplementary material for "High radiosensitivity in the conifer Norway spruce (*Picea abies*) due to less comprehensive mobilisation of protection and repair responses compared to the radiotolerant *Arabidopsis thaliana*": Table legends for supplemental data

**Table S1**. Table of experiments with gamma irradiation of *A. thaliana* and Norway spruce from day 7 after sowing.

**Table S2.** Primers for q-PCR analysis of DNA repair-, cell cycle-, antioxidant- and retrograde signaling-related genes after gamma irradiation of *A. thaliana* and Norway spruce.

**Table S3.** Differentially expressed genes in DNA damage repair (DDR) pathways in *A. thaliana* and Norway spruce shoots after 48 h of gamma irradiation from day 7 after sowing compared to unexposed controls. Red = upregulation, blue = downregulation, black rectangles = significant difference from unexposed control (p > 0.05). The fold change 0 = no significant difference in expression. HR = Homologous recombination, NHEJ = Non homologous end joining.

**Table S4.** Differentially expressed cell cycle-related genes in *A. thaliana* and Norway spruce shoots after 48 h of gamma irradiation from day 7 after sowing compared to unexposed controls. Red = upregulation, blue = downregulation, black rectangles = significant difference from unexposed control (p > 0.05). The fold change 0 = no significant difference in expression.

**Table S5.** Differentially expressed endoreduplication-related genes pathway in *A. thaliana* and Norway spruce shoots after 48 h of gamma irradiation compared to the unexposed controls. Red = upregulation, blue = downregulation, black rectangles = significant difference from unexposed control (p > 0.05). The fold change 0 = no significant difference in expression.

**Table S6**. Differentially expressed genes related to histone-/DNA modification and epigenetic regulation in *A. thaliana* and Norway spruce shoots after 48 h of gamma irradiation from day 7 after sowing compared to unexposed controls. Red = upregulation, blue = downregulation, black rectangles = significant difference from unexposed control (p > 0.05). The fold change 0 = no significant difference in expression.

**Table S7.** Differentially expressed genes related to hormone biosynthesis and signalling pathways in *A. thaliana* and Norway spruce shoots after 48 h of gamma irradiation from day 7 after sowing compared to unexposed controls. Red = upregulation, blue = downregulation, black rectangles = significant difference from unexposed control (p > 0.05). The fold change 0 = no significant difference in expression.

**Table S8.** Differentially expressed genes in antioxidant biosynthesis and signalling pathways in *A. thaliana* and Norway spruce shoots after 48 h of gamma irradiation from day 7 after sowing compared to unexposed controls. Red = upregulation, blue = downregulation, black rectangles = significant difference from unexposed control (p > 0.05). The fold change 0 = no significant difference in expression.

**Table S9.** Differentially expressed genes related to cell wall biogenesis and maintenance resulting in *A. thaliana* and Norway spruce shoots after 48 h of gamma irradiation from day 7 after sowing compared to unexposed controls. Red = upregulation, blue = downregulation, black rectangles = significant difference from unexposed control (p > 0.05). The fold change 0 = no significant difference in expression.

**Table S10.** Differentially expressed photosynthesis-related genes in *A. thaliana* and Norway spruce shoots after 48 h of gamma irradiation from day 7 after sowing compared to unexposed controls. Red = upregulation, blue = downregulation, black rectangles = significant difference from unexposed control (p > 0.05). The fold change 0 = no significant difference in expression.

**Table S11.** Differentially expressed energy metabolism-related genes in *A. thaliana* and Norway spruce shoots after 48 h of gamma irradiation from day 7 after sowing compared to unexposed controls. Red = upregulation, blue = downregulation, black rectangles = significant difference from unexposed control (p > 0.05). The fold change 0 = no significant difference in expression.

**Table S12.** Gene names and gene function. Full gene names and their functions based on information available in the databases TAIR (www.arabidopsis.org) and RARGE (rarge.gsc.riken.jp)
