## Supplementary material for "High radiosensitivity in the conifer Norway spruce (*Picea abies*) due to less comprehensive mobilisation of protection and repair responses compared to the radiotolerant *Arabidopsis thaliana*": Table S1

| Experiments | Duration of irradiation (h) | Dose rates (mGy h^-1^) | Time of sampling/ measurements | Biological replicates /per dose rate | No. of seedlings |
| --- | --- | --- | --- | --- | --- |
| 1. Growth studies |  |  |  |  |  |
| A. At the end of the irradiation | 48 | 0-290* | 48 h |  | *A. thaliana*-29-40, *P. abies*-28-35. |
| B. Post-irradiation period in *A. thaliana* |  |  | Up to 77 days |  | 23-24 seedlings |
| C. Post irradiation period in *P. abies* |  |  | Up to 59 or 77 days |  | 11 to 22 seedlings |
| 2. Histology and cytology | 48 | 0-290 | 48 h |  | 5 plants per dose rate, for both species. |
| 3. Comet assays |  |  |  |  |  |
| A. At the end of irradiation | 48 | 0-290 | 48 h | 3 | *A. thaliana* - 5 plants  *P. abies* - 3 plants, in each biological replicate. |
| B. Post-irradiation |  |  | 77 days |  |  |
| 4. RNA-seq | 48 | For *A. thaliana* 0-290, for *P. abies* 0-100 | 48 h | 4 | *A. thaliana* -20 - 30 (shoots only), *P. abies* – 4 (shoots only), in each replicate. |
| 5. Q-PCR | 12, 24, 48 | 0-100 | 12, 24, 48 h | 4 | *A. thaliana* – 30 (shoots only), *P. abies* - 8 (shoots only) in each replicate. |
| 6. Antioxidant assays | 12, 24, 48 | 0-100 | 12, 24, 48 h | 3-4 | *A. thaliana*- 50-100 mg/ sample, *P. abies*- 50-100 mg/sample, in each replicate. |

*In one of the two repeated experiment the highest dose rate obtained was 250 mGy h^-1^, due to the half life of the Co^60^ core.
